# Mechanism, and treatment of anti-CV2/CRMP5 autoimmune pain

**DOI:** 10.1101/2024.05.04.592533

**Authors:** Laurent Martin, Harrison J. Stratton, Kimberly Gomez, Le Duy Do, Santiago Loya-Lopez, Cheng Tang, Aida Calderon-Rivera, Dongzhi Ran, Venkatrao Nunna, Shreya S. Bellampalli, Liberty François-Moutal, Nicolas Dumaire, Lyuba Salih, Shizhen Luo, Frank Porreca, Mohab Ibrahim, Véronique Rogemond, Jérôme Honnorat, Rajesh Khanna, Aubin Moutal

**Author notes:** To whom correspondence should be addressed: Dr. Aubin Moutal, Saint Louis University - School of Medicine, Department of Pharmacology and Physiology, 1402 S. Grand Blvd., Schwitalla Hall, Room 432, Saint Louis, MO 63104, USA. Co-first authors (Laurent Martin, Harrison J. Stratton). Contributed equally to this work (Rajesh Khanna, Aubin Moutal).

## Abstract

Paraneoplastic neurological syndromes arise from autoimmune reactions against nervous system antigens due to a maladaptive immune response to a peripheral cancer. Patients with small cell lung carcinoma or malignant thymoma can develop an autoimmune response against the CV2/collapsin response mediator protein 5 (CRMP5) antigen. For reasons that are not understood, approximately 80% of patients experience painful neuropathies. Here, we investigated the mechanisms underlying anti-CV2/CRMP5 autoantibodies (CV2/CRMP5-Abs)-related pain. We found that patient-derived CV2/CRMP5-Abs can bind to their target in rodent dorsal root ganglia (DRG) and superficial laminae of the spinal cord. CV2/CRMP5-Abs induced DRG neuron hyperexcitability and mechanical hypersensitivity in rats that were abolished by preventing binding to their cognate autoantigen CRMP5. The effect of CV2/CRMP5-Abs on sensory neuron hyperexcitability and mechanical hypersensitivity observed in patients was recapitulated in rats using genetic immunization providing an approach to rapidly identify possible therapeutic choices for treating autoantibody-induced pain including the repurposing of a monoclonal anti-CD20 antibody that selectively deplete B-lymphocytes. These data reveal a previously unknown neuronal mechanism of neuropathic pain in patients with paraneoplastic neurological syndromes resulting directly from CV2/CRMP5-Abs-induced nociceptor excitability. CV2/CRMP5-Abs directly sensitize pain responses by increasing sensory neuron excitability and strategies aiming at either blocking or reducing CV2/CRMP5-Abs can treat pain as a comorbidity in patients with paraneoplastic neurological syndromes.

## Introduction

Paraneoplastic neurological syndromes manifest as autoimmune reactions against nervous system antigens triggered by a maladaptive immune response to a peripheral cancer (1). Patients with small cell lung carcinoma or malignant thymoma can develop an autoimmune paraneoplastic syndrome with anti-CV2/collapsin response mediator protein 5 (CRMP5) autoantibodies (CV2/CRMP5-Abs) (2–4). Among other symptoms such as encephalitis, myelopathy or cerebellar ataxia, 80% of patients with CV2/CRMP5-Abs also experience idiopathic painful neuropathy, often manifesting as symmetric or asymmetric polyradiculoneuropathy (2). Pain frequently serves as the initial symptom that prompts patients to seek medical attention, ultimately leading to the diagnosis of their autoimmune disease (5). The remarkably high prevalence of pain in this patient population surpasses that of many other well-characterized pain-related diseases (6–8).

Our goal was to understand how anti-CV2/CRMP5 autoimmunity causes neuropathic pain. Additionally, we aimed to develop a new model of anti-CV2/CRMP5 painful neuropathy, reproducing the pain symptoms observed in patients allowing mechanistic investigation and assessment of possible therapeutic interventions. Human CV2/CRMP5-Abs have been shown to bind to human dorsal root ganglia (DRG) where sensory neurons are located (4, 5). CRMP5 is the most distant member (<50% homology) of the collapsin response mediator protein family (9, 10) and is highly expressed in the central nervous system during embryonic development (11) where it regulates dendritic growth and Purkinje cell maturation (12, 13). After birth, CRMP5 expression is downregulated but retained only in the midbrain (11), DRG and spinal cord (10). While patients with anti- CV2/CRMP5 paraneoplastic neurological syndrome commonly present primarily with pain, the possible contribution of CRMP5 expression or function to pain has not been previously explored.

Immunotherapy aimed at reducing autoantibody levels has been reported to reduce pain reported by patients (2). Therefore, we postulated that CV2/CRMP5-Abs might directly sensitize sensory neurons leading to promote pain. Our investigation revealed that patient-derived CV2/CRMP5-Abs induced a hyperexcitability phenotype in cultured rodent DRG sensory neurons. Remarkably, this hyperexcitability correlated with heightened sensitivity to mechanical stimuli in rats. By immunizing rats against CRMP5, we successfully replicated the pain phenotype observed in patients with CV2/CRMP5-Abs. Leveraging this novel model, we assessed the effectiveness of common painkillers and a monoclonal antibody, potentially offering valuable treatment options for patients experiencing autoantibody-induced pain.

## Results

### Serum from patients with CV2/CRMP5-Abs stain their target CRMP5 in dorsal root ganglia and spinal cord

The concomitant occurrence of CV2/CRMP5-Abs and pain symptoms in the early stages suggests that these autoantibodies likely play a causal role in the pain experienced by patients (5). To directly explore the relationship between CV2/CRMP5-Abs and pain, we initially investigated whether sera from patients could identify their target along the pain neuraxis. Previous studies have reported positive autoimmuno-reactivity of anti-CV2/CRMP5 autoantibodies in various brain regions, including the brainstem and the cerebellum (14, 15). Additionally, there is evidence that these autoantibodies can stain the human sciatic nerve and the dorsal root ganglia (DRG) (4, 5). We assembled a collection of sera from 22 patients diagnosed with both anti-CV2 autoantibodies and neuropathy (**Table 1**). Our cohort included samples from 16 males and 6 females, collected during diagnosis and preserved for research purposes. No apparent sex-based prevalence associated with the development of CV2/CRMP5-Abs was reported (2, 15).

**Table 1:** Anti-CV2/CRMP5 positive sera used in this study. . Sex, origin of the primary tumor and age at diagnostics are indicated. All patients had a type of neuropathy and those who reported pain are indicated.

| Patient # | Sex | Type of neuropathy | Pain | Primary tumor | Age at diagnostic |
| --- | --- | --- | --- | --- | --- |
| 1 | M | Motor and sensory neuropathy | Yes | Small cell lung cancer | 56 |
| 2 | M | Sub-acute sensory neuropathy | Yes | Small cell lung cancer | 75 |
| 3 | M | Motor and sensory neuropathy | Yes | Small cell lung cancer | 60 |
| 4 | F | Peripheral neuropathy | Yes | Small cell lung cancer | 70 |
| 5 | F | Peripheral neuropathy | Yes | Small cell lung cancer | 70 |
| 6 | F | Sensory neuropathy and Lambert-Eaton syndrome | Yes | Small cell lung cancer | 64 |
| 7 | F | Sensory neuropathy and Lambert-Eaton syndrome | Yes | Small cell lung cancer | 64 |
| 8 | M | Peripheral neuropathy | Yes | NA | 70 |
| 9 | M | Sensory neuropathy (and cerebellar syndrome) | Yes | Small cell lung cancer | 64 |
| 10 | M | Motor and sensory neuropathy | Yes | Bladder | 54 |
| 11 | F | Motor and sensory neuropathy | Yes | ? | 55 |
| 12 | M | Lambert-Eaton syndrome | Yes | Small cell lung cancer | 78 |
| 13 | F | Polyneuropathy | Yes | ? | 56 |
| 14 | M | Sensory neuropathy | Yes | Non small cell lung cancer | 76 |
| 15 | M | Sensory neuropathy | Yes | ? | 62 |
| 16 | M | cerebellar syndrome | No | Small cell lung cancer | 63 |
| 17 | M | Sensory neuropathy | No | Small cell lung cancer | 56 |
| 18 | M | Sensory neuropathy | No | Small cell lung cancer | 80 |
| 19 | M | uveitis | No | Small cell lung cancer | 62 |
| 20 | M | encephalitis | No | intestinal | 69 |
| 21 | M | Sensory neuropathy | No | Small cell lung cancer | 45 |
| 22 | M | Motor and sensory neuropathy | No | Small cell lung cancer | 60 |

We found that sera from patients exhibited positive staining in the neuronal somata of rat DRG neurons (**Figure 1A**). Additionally, strong labeling was observed in the spinal dorsal horn, where nociceptive neurons synapse onto second-order neurons (**Figure 1A and S1**). This signal was specifically localized to laminae I and IIo of the dorsal spinal cord, suggesting that CV2/CRMP5-Abs may identify nociceptive fibers (16). To further investigate this immunoreactivity, we conducted an additional co-staining using a validated anti-CRMP5 antibody (17) (**Figure S1**). While a control serum from an CV2/CRMP5-Abs-negative patient showed no staining, we observed comparable staining patterns between CV2/CRMP5-Abs and CRMP5 in the rat DRG and spinal dorsal horn strongly suggesting that observed immunoreactivity is related to CRMP5. We additionally confirmed that CRMP5 is expressed in human DRG neurons (**Figure S2**).

**Figure 1:**
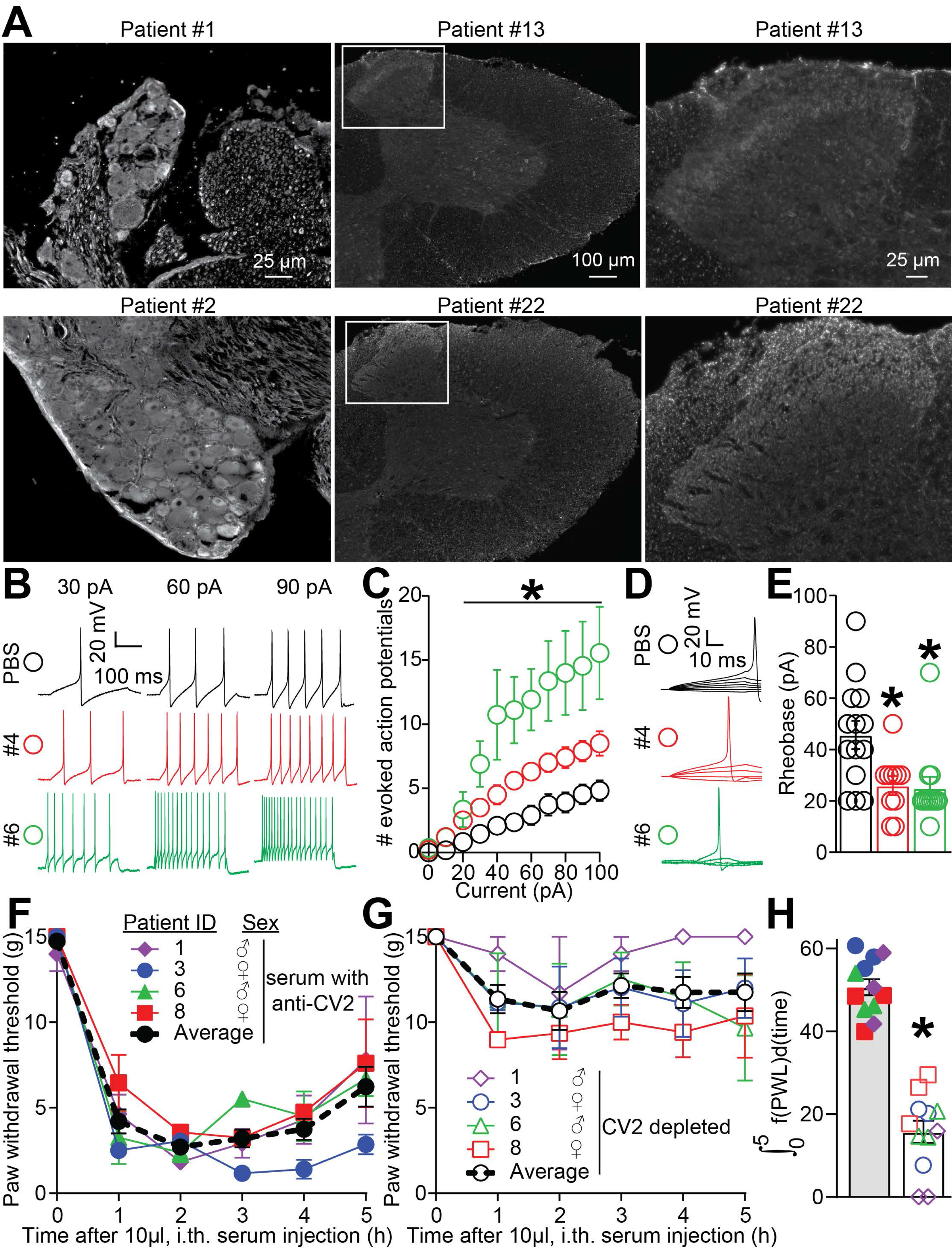
CV2/CRMP5-Abs label nociceptive structures and induce mechanical hypersensitivity and DRG neuron hyperexcitability. (**A**) Micrographs of rat dorsal root ganglia (DRG) and spinal cord immunolabelled with anti-CV2 sera from three patients. CV2/CRMP5-Abs positively labelled neuronal soma in DRG and superficial laminae in the dorsal horn of the lumbar spinal cord. The right panels show a magnification of boxed regions of spinal cord sections. (**B**) Representative recordings of evoked action potentials recorded from small-diameter DRG neurons in response to depolarizing current injection of 30, 60, and 90 picoamperes (pA). Female rat DRG neurons were treated overnight with serum from patient #4 or #6 (1/100 dilution). (**C**) Quantification of the number of evoked action potentials in response to 0-100 pA of injected current. *p<0.05, multiple Mann-Whitney tests. Representative traces (**D**) and (**E**) bar graph with scatter plot showing a decreased rheobase in cells treated with the serum from patient #4 or #6. n=11 cells per condition; error bars indicate mean ± SEM, *p<0.05, Mann-Whitney test. (**F**) Graph showing the paw withdrawal threshold of male rats injected intrathecally (i.th.) with 10 µL of the indicated positive CV2/CRMP5-Abs or (**G**) depleted (cross-adsorbed with purified CRMP5) sera. CV2/CRMP5-Abs containing sera consistently elicited mechanical hypersensitivity while depleted sera had no significant effect. n = 12; 3 rats per CV2/CRMP5-Abs serum; 4 different CV2/CRMP5-Abs sera. The black line shows the average of all patients tested. Error bars indicate mean ± SEM, *p<0.05, two-way ANOVA. (**H**) Bar graph with scatter plot showing the area under the curve for the data in **F** and **G**, *p<0.05, Mann-Whitney test.

The results above raise questions about how these autoantibodies may interact with neurons. To address this, we employed a technique called proximity ligation assay (PLA), which permits detection of protein-protein interactions in situ at endogenous protein levels (25). In rat sensory neurons exposed overnight to CV2/CRMP5- Abs, we observed positive human IgG/CRMP5 PLA signals only when we permeabilized fixed cells prior to immunostaining. This indicates that human autoantibodies can bind to intracellular autoantigens, at least within sensory neurons (**Figure S3**). As a negative control, we also utilized sera from the same patients, depleted by pre-adsorption on magnetic beads coated with purified CRMP5 (**Figure S3**). Additionally, we fractionated DRG neurons treated with patient serum to isolate cytosolic, membrane, nuclear, and cytoskeletal fractions. Notably, human autoantibodies were detected in the cytosol of neurons through immunoblotting (**Figure S3**). Altogether, these findings highlight that anti-CV2 autoantibodies can specifically identify their target autoantigen within the cytoplasm of DRG sensory neurons and spinal cord dorsal horn neurons.

We initially explored whether CV2/CRMP5-Abs could alter the overall function of DRG neurons. Employing a functional fingerprinting technique called constellation pharmacology, we used live cell calcium imaging to detect changes directly related to the sensitization of pain-relevant receptors and ion channels (18, 19). This approach uses several agonists to ligand gated ion channels (AITC: TRPA1, noxious cold; ATP: P2X and P2Y, inflammation; menthol: TRPM8, innocuous cold; capsaicin: TRPV1, noxious heat) and G-coupled protein receptors (acetylcholine: nAChR; histamine: histamine receptors, itch) to identify functional sensory neuron populations involved in heat, cold, itch or inflammatory pain (19). As a reference control, we compared the serum of an CV2/CRMP5-Abs-negative patient with sera from patients #3 and #6 (**Figure S4A**). Surprisingly, CV2/CRMP5-Abs exposure overnight had no effect on the proportion of DRG neurons responding to more than one receptor agonist (**Figure S4B**) or to each receptor agonist (**Figure S4C**). Additionally, the peak calcium response to each compound remained unchanged (**Figure S3D**). We further analyzed the peak response to a depolarizing stimulus within each functional sensory neuron subpopulation and found no alteration due to CV2/CRMP5-Abs exposure (**Figure S4E**). Finally, assessing the size of DRG neurons in relation to their functional response revealed no significant difference (**Figure S4F**).

### CV2/CRMP5-Abs increase sensory neuron excitability and sensitize mechanical responses in rats

Because CV2/CRMP5-Abs label sensory neurons in the DRG and are associated with painful neuropathy in humans, we next investigated whether these autoantibodies could cause hyperexcitability of DRG neurons, a crucial feature of pain sensitization in both rodents and humans (20–22). To explore this, we cultured rat DRG neurons and exposed them to CV2/CRMP5-Abs containing serum overnight. Using patch clamp electrophysiology, we recorded excitability by evoking action potentials through stepwise current injections. DRG neurons treated with CV2/CRMP5-Abs exhibited significantly higher evoked action potentials compared to control neurons, starting at 30 pA (**Figure 1B-C**). Moreover, the rheobase, which represents the current needed to elicit a single action potential, was lower in CV2/CRMP5-Abs-treated neurons (**Figure 1D-E**). In summary, our data strongly suggests that CV2/CRMP5-Abs enhance the excitability of sensory neurons, potentially contributing to pain hypersensitivity.

We next asked if anti-CV2/CRMP5 autoantibodies could induce pain-like behaviors in rodents. Specifically, we focused on mechanical hypersensitivity, a common symptom observed in patients with neuropathic pain (23, 24). To assess this, we employed an intrathecal catheter to inject rats with patients’ serum containing CV2/CRMP5-Abs, comparing it to antibody-depleted controls. We then used Von Frey filaments to measure mechanical sensitivity on the hindpaw. Remarkably, all four sera tested lowered mechanical thresholds for at least 5 hours, while antibody depletion effectively prevented this effect (**Figure 1F-H**). These findings demonstrate that CV2/CRMP5-Abs can sensitize the pain pathway, leading to the manifestation of mechanical hypersensitivity.

### CV2/CRMP5-Abs bind to surface accessible epitopes on CRMP5

While our findings (**Figure 1)** are compelling, they do not definitively establish a causal link between CV2/CRMP5-Abs and neuronal sensitization or pain responses via their antigen binding fragment (Fab). We used a peptide array approach to map the protein sequence of CRMP5 using 15-mer peptides with 3 amino acid increments. We consistently identified three distinct epitopes (Peptides 53, 94, and 146) (**Figure 2A**), along with Peptide 142 that were recognized by sera from six of seven patients (sequences shown in **Figure S5**). Notably, patient characteristics (age, sex, neuropathy type, and primary tumor) did not significantly impact epitope recognition (**Table 1 and Figure 2A**). These four epitopes, uniquely associated with CRMP5 (PDB ID: 4B90, (26)), were exposed on the protein’s surface and accessible for binding by CV2/CRMP5-Abs under native conditions (**Figure S5B**). We next utilized epitope peptides to mask the Fab domain of CV2/CRMP5-Abs. Peptides 53, 142, and 146 were selected for these experiments while peptide 94 was excluded due to its solubility limitations. Employing an ELISA approach, we successfully blocked the Fab region of human autoantibodies using these epitope peptides (100 ng/ml each). This prevented their binding to the target protein CRMP5 (**Figure 2B**). Intriguingly, purified CRMP5 at the same concentration exhibited a similar level of inhibition, with less than 25% residual binding (**Figure 2B**). Furthermore, we confirmed that the epitope peptides 53, 142, and 146 effectively blocked CV2/CRMP5-Abs immunoreactivity to DRG neurons (**Figure 2C**). These findings demonstrate that CV2/CRMP5-Abs specifically bind to the peptide sequences 53, 142, and 146 on their target, CRMP5. Consequently, these peptides can be employed to prevent the binding of autoantibodies to their endogenous target.

**Figure 2:**
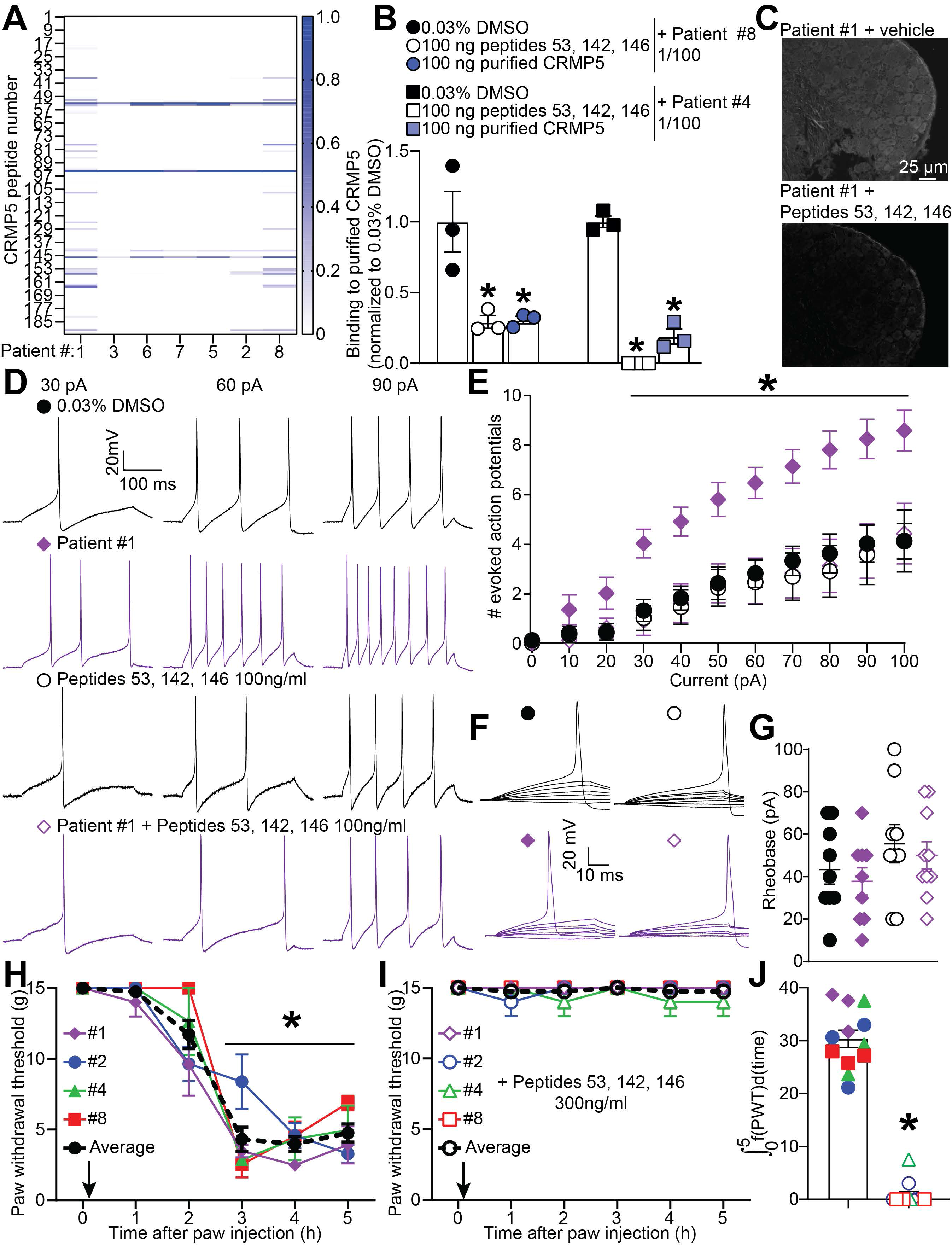
Blocking CV2/CRMP5-Abs with epitope peptides prevents the sensitization of sensory neurons and mechanical hypersensitivity in rats. (**A**) Heatmap of the immunoreactivity of CV2/CRMP5-Abs sera hybridized on a CRMP5 peptide array mapping the entire sequence of the protein in 15-mer peptides with 3 amino acid increments. Four main epitopes on CRMP5 are targeted by the CV2/CRMP5-Abs. (**B**) Bar graph with scatter plot showing that peptides 53, 142 and 146 can block the binding of CV2/CRMP5-Abs from patients #8 and #4 to purified CRMP5. 0.03% DMSO is the vehicle, purified CRMP5 was used as a positive control to achieve maximal displacement of the CV2/CRMP5-Abs. n=3 independent measures from an average of 3 repeats. (**C**) Micrograph of a rat DRG immunolabelled with CV2/CRMP5-Abs positive serum (1/100) from patient #1 and then with blocking peptides 53, 142, 146. Blocking peptides abolished the immunoreactivity of CV2/CRMP5-Abs for their protein target CRMP5. (**D**) Representative recordings of evoked action potentials recorded from small-diameter DRG neurons treated with serum from patient #1 (1/100 dilution) in combination with 100 ng/ml of peptides 53, 142 and 146 as indicated, overnight in response to depolarizing current injection of 30, 60, and 90 pA. (**E**) Quantification of the number evoked action potentials in response to 0-100 pA of injected current. n=9- 10 cells per condition, *p<0.05, multiple Mann-Whitney tests. (**F**) traces and (**G**) bar graph with scatter plot showing unchanged rheobase in cells treated with the serum from patient #1 with blocking peptides as indicated. Error bars indicate mean ± SEM, *p<0.05, Mann-Whitney test. (**H**) Graph showing the paw withdrawal threshold of rats injected with 15 µl of the indicated CV2/CRMP5-Abs positive sera (1/10 dilution) or with (**I**) blocking peptides 53, 142 and 146 (300 ng/ml) in the paw. Anti-CV2 containing sera elicited mechanical hypersensitivity which was prevented by blocking the CV2/CRMP5-Abs with their epitope peptides. Error bars indicate mean ± SEM, *p<0.05, two-way ANOVA. (**J**) bar graph with scatter plot showing the area under the curve of the data in G and H and color coded per treatment groups (n=3 each), error bars indicate mean ± SEM, *p<0.05, Mann- Whitney. Experimenters were blind to the treatment groups.

### Blocking CV2/CRMP5-Abs prevents DRG neuron excitability and mechanical hypersensitivity

To explore whether the increased excitability of DRG neurons exposed to CV2/CRMP5-Abs (**Figure 1D**) was solely due to the action of these autoantibodies, we employed peptides 53, 142, and 146 to block the Fab region. We treated cultured rat DRG neurons with whole serum (diluted 1/100) from patient #1 and added 100 ng/ml of the aforementioned peptides (**Figure 2D**). Consistent with our previous observations (**Figure 1C**), CV2/CRMP5- Abs increased sensory neuron firing frequency (**Figure 2D**). Peptides 53, 142, and 146 (at 100 ng/ml) alone had no impact on action potential firing. However, sensory neuron hyperexcitability (**Figure 2D-E**) was successfully prevented by blocking the binding of autoantibodies using these peptides. Rheobase remained unaffected (**Figure 2F-G**).

Next, we replicated this approach in vivo to investigate whether mechanical hypersensitivity induced by treatment with CV2/CRMP5-Abs was exclusively mediated by the Fab region of autoantibodies. We directly injected anti- CRMP5 sera from four patients (diluted 1/10) into the paw of rats, allowing direct exposure of nociceptive terminals in the skin to the autoantibodies. Remarkably, hindpaw injection of CV2/CRMP5-Abs from these patients led to robust mechanical hypersensitivity (**Figure 2H**), demonstrating that autoantibodies can sensitize pain responses when present in the skin. Crucially, blocking the Fab region of human anti-CRMP5 using peptides 53, 142, and 146 prevented the development of mechanical hypersensitivity for all anti-CV2 sera tested (**Figure 2I-J**). Collectively, our results underscore that CV2/CRMP5-Abs-induced mechanical hypersensitivity is mediated by the binding of autoantibodies to their target protein, CRMP5.

### Preclinical replication of CV2/CRMP5 autoimmune pain

After establishing that autoantibodies play a pivotal role in sensory neuron sensitization and mechanical hypersensitivity in anti-CV2/CRMP5 autoimmune neuropathy, our next goal was to determine if paraneoplastic neurological syndromes could be replicated preclinically allowing for further mechanistic investigation and determination of potential therapeutic strategies. Interestingly, direct infusion of patient autoantibodies into animals often failed to fully replicate the symptoms seen in paraneoplastic neurological syndromes (27, 28). To overcome this challenge, we adopted an alternative approach: DNA immunization (29). By injecting a plasmid containing the CRMP5 coding sequence, we invoked muscle cells to produce the antigen, which was then recognized by the immune system. This immunization technique activated both T- and B-lymphocytes, leading to the production of CRMP5 autoantibodies in rats (**Figure S6A**).

Our peptide array analysis revealed that, akin to humans, anti-CRMP5 autoantibodies from rat serum specifically targeted two epitopes: Peptides 53 and 146 (**Figure S6B**). Notably, rats immunized against CRMP5 concurrently developed bilateral mechanical hypersensitivity as their autoantibody levels increased in the serum (**Figure 3A- B** and **S6A**). Surprisingly, we did not observe thermal hypersensitivity (**Figure 3C-D**), suggesting that CRMP5 autoimmunity predominantly sensitizes mechanical responses. To further explore this link between mechanical hypersensitivity and pain, we employed the mechanical conflict avoidance assay. In this test, rats had to navigate a field of sharp probes to escape a bright light aversive stimulus and reach a safe, dark chamber (30). Strikingly, rats with CRMP5 autoimmunity took longer to cross the probe-filled field compared to control rats, indicating that their heightened tactile sensitivity was associated with an aversive (painful) effect (**Figure S6C**). Consistent with the lack of sex differences in pain reported clinically, these findings held true for both male (**Figure 3A-D**) and female rats (**Figure 3E-H**).

**Figure 3:**
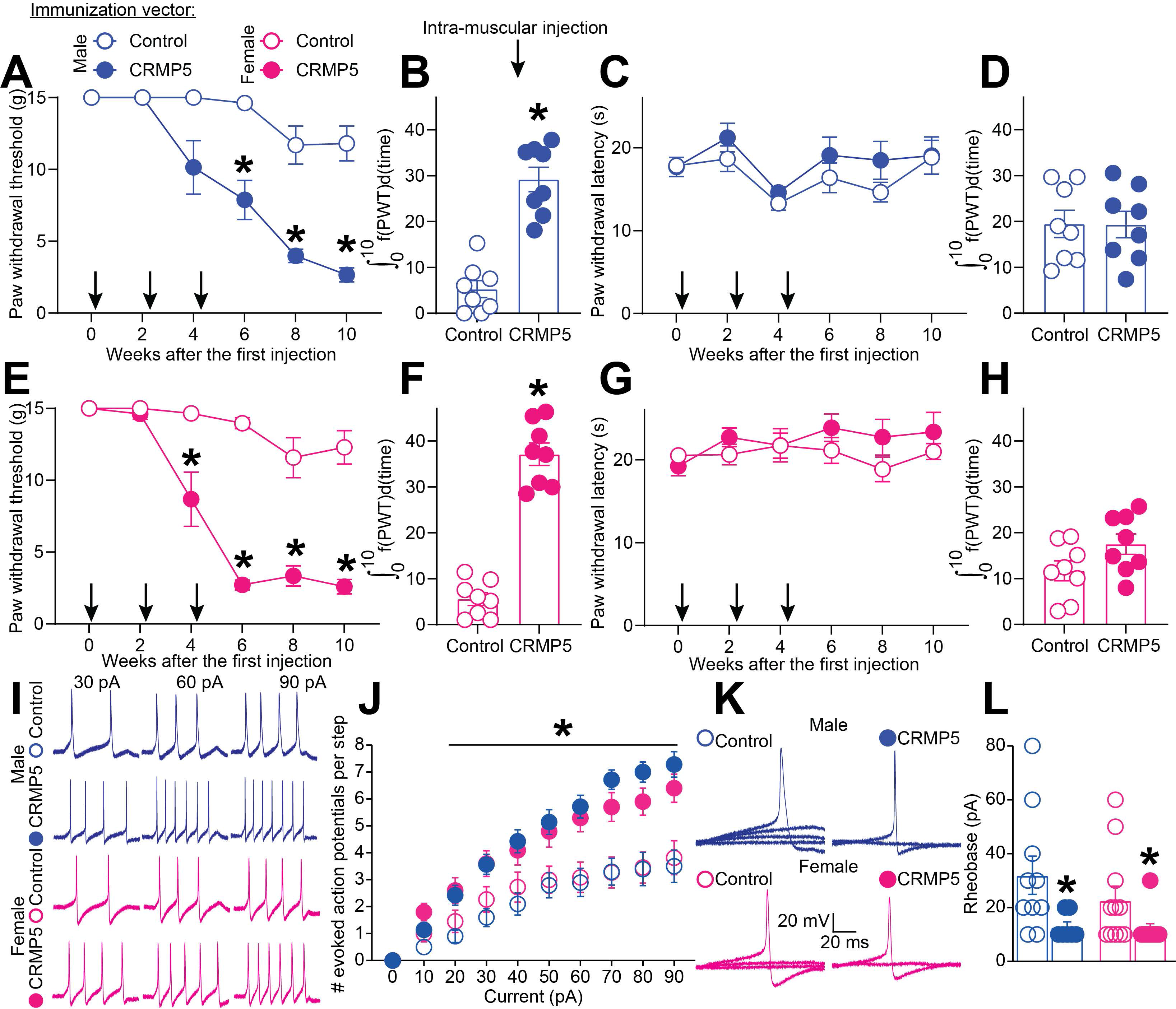
CRMP5 auto-immunity induces mechanical allodynia and hyperexcitability in both male and female rats. Rats were immunized by 3 injections (indicated by arrows) of 50 µg of pCMV2 plasmid allowing for the expression of CRMP5 or control (empty) in the spinodeltoidus muscle. Rats received an intramuscular injection of a plasmid (pCMV2-CRMP5) carrying the coding sequence for CRMP5 on day 0 and then 2 booster shots at weeks 2 and 4 after the first injection. Graph showing the paw withdrawal threshold of (**A**) male and (**E**) female rats injected with the CRMP5 coding plasmid compared to the control empty plasmid (n=8 animals per group; *p<0.05, two-way ANOVA). In both sexes, rats injected with the CRMP5 coding plasmid developed mechanical hypersensitivity. Bar graph with scatter plot showing the area under the curve for the mechanical thresholds in (**B**) male and (**F**) female rats injected as described above (n=8 animals per group; *p<0.05, Mann-Whitney test). Rats were tested for their thermal thresholds using the Hargreave’s test and no difference was found in (**C**) male and (**G**) female rats injected with the CRMP5 coding plasmid compared to the control empty plasmid (n=8 animals per group; *p<0.05, two-way ANOVA). Bar graph with scatter plot showing the area under the curve for the thermal thresholds in (**D**) male and (**H**) female rats injected as described above (n=8 animals per group; *p<0.05, Mann-Whitney test. Error bars represent mean ± SEM. (**I**) Representative recordings in response to a depolarizing current step to evoke action potentials (APs) in sensory neurons from male and female rats injected with plasmid expressing CRMP5 or a control plasmid. (**J**) Summary of the number of APs in the indicated conditions (n=10 cells per condition). (**K**) Representative recordings in response to various steps of depolarizing current to measure rheobase in sensory neurons prepared from rats with CRMP5 autoimmunity or control. (**L**) Summary of the measured rheobase in indicated conditions (n=10 each). Asterisks indicate significance compared with control *p<0.05, Mann-Whitney test. Error bars represent mean ± SEM.

Subsequently, we cultured male and female DRG neurons and recorded their action potential firing (**Figure 3I**). Neurons from rats immunized against CRMP5 exhibited an increased frequency of evoked action potential discharge compared to control neurons (**Figure 3J**). Additionally, the rheobase (the minimum current required to elicit an action potential) was lower in CRMP5-immunized rats (**Figure 3K-L**). Collectively, these results strongly suggest that primary afferent neurons become sensitized in the context of CRMP5 autoimmunity.

To further validate our approach, we conducted an ex vivo assay. By applying a depolarization stimulus, we induced the release of calcitonin gene-related peptide (CGRP) from the spinal cord, which we quantified using ELISA. Remarkably, this assay demonstrated increased CGRP release in rats with autoimmunity against CRMP5, highlighting the association with pain (**Figure S6D**). Altogether, our findings underscore the successful use of DNA immunization to induce CRMP5 autoimmunity, faithfully recapitulating the pain symptoms and autoantibody profiles observed in patients.

### Preclinical evaluation of potential therapies for CRMP5 autoimmune painful neuropathies

High-dose IV corticosteroids are commonly used to manage symptoms associated with CRMP5 autoimmunity including pain (2). While this therapy can be effective, it is associated with a multitude of undesired side effects. For this reason, we evaluated the efficacy of commonly used clinical therapies including acetaminophen, ibuprofen (a non-steroidal anti-inflammatory drug), amitriptyline and duloxetine (reuptake blockers), gabapentin and morphine (31) on mechanical allodynia in rats immunized with the CRMP5 coding sequence. Since male and female rats exhibited similar behavior profiles over time, we conducted drug tests on only one sex (**Figure 3**). Consistent with clinical observations, pain induced by CRMP5 autoimmunity was reversed by ibuprofen (100 mg/kg, p.o.), amitriptyline (50 mg/kg, p.o.) or duloxetine (30 mg/kg, p.o.) (**Figure S7A**) but not acetaminophen (200 mg/kg, p.o.). As anticipated (2), morphine effectively alleviated mechanical hypersensitivity in rats with CRMP5 autoimmunity (**Figure S7A**). Interestingly, gabapentin (30 mg/kg, p.o.) had little to no effect (**Figure S7A**). Overall, amitriptyline, duloxetine, and morphine demonstrated the strongest impact on CRMP5 autoimmunity-induced pain (**Figure S7B**).

These results highlight the effectiveness of clinically used drugs in managing pain for patients with CRMP5 autoimmune neuropathy. However, these drugs typically offer short-term relief based on their pharmacokinetic profiles and do not represent long-term treatment options for this patient population. To explore a more lasting solution, we investigated the benefits of an anti-CD20 monoclonal antibody (Genentech, 4mg/kg, i.p. (32)) for its potential as a long-term therapy to reverse pain from CRMP5 autoimmune painful neuropathy. Anti-CD20 specifically targets B lymphocytes as autoantibody producing cells (33) and is currently used to treat conditions such as non-Hodgkin lymphoma, chronic lymphocytic leukemia, and refractory rheumatoid arthritis. We administered anti-CD20 on day 56 and then day 63 to rats with CRMP5 autoimmunity. The treatment effectively depleted B lymphocytes present in their serum (**Figure S8A-B**) and lowered serum levels of anti-CRMP5 autoantibodies (**Figure S8C**). During our injection protocol, we measured mechanical withdrawal thresholds (**Figure 4A**) and observed that Anti-CD20 completely reversed the mechanical hypersensitivity induced by CRMP5 autoimmunity (**Figure 4B**). Since CRMP5 autoimmunity leads to hyperexcitability of DRG sensory neurons, we further investigated whether anti-CD20 could reverse this sensitization state (**Figure 4C**). Remarkably, the excitability profile of DRG neurons prepared from rats with CRMP5 autoimmunity and treated with anti-CD20 was comparable to control levels (**Figure 4D**). Additionally, the rheobase of neurons from anti- CD20-treated rats returned to the level of control rats (**Figure S8D-E**). Serum cytokine profiling revealed that anti-CD20 treatment normalized most dysregulated cytokines in rats with CRMP5 autoimmunity (**Figure S8F**). These findings demonstrate that anti-CD20 has the potential to fully reverse sensory neuron sensitization and pain induced by CRMP5 autoimmunity, offering hope for treating CRMP5 painful autoimmune neuropathy in humans.

**Figure 4:**
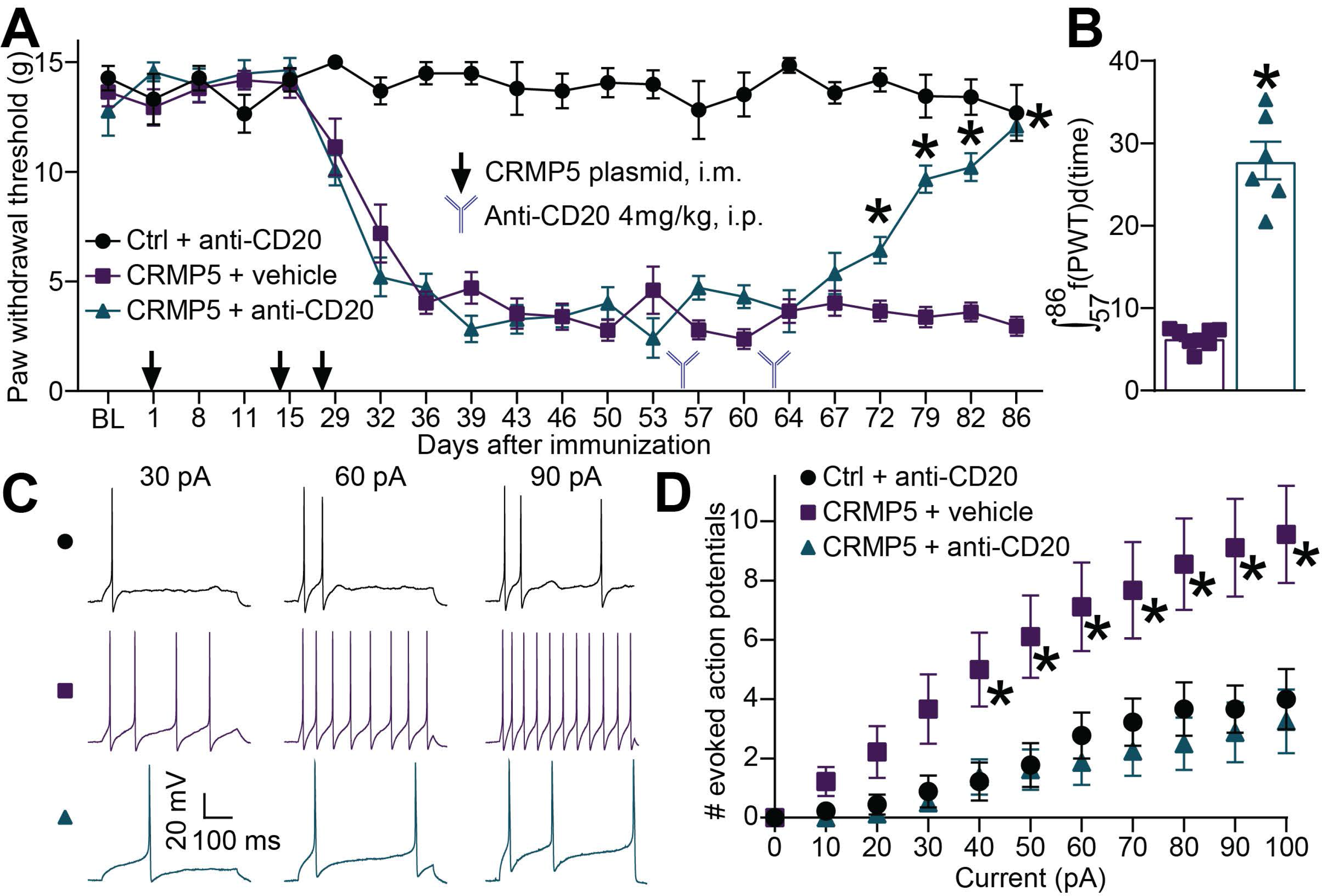
The anti-CD20 monoclonal antibody reverses CRMP5 autoimmunity induced mechanical hypersensitivity and hyperexcitability. Rats were immunized against CRMP5 and developed stable mechanical hypersensitivity up to day 86 after the first intramuscular injection. (**A**) Graph showing the paw withdrawal threshold of rats over time. Black arrows show the 3 plasmid injections necessary for the induction of the model. Anti-CD20 injection (4 mg/kg, i.p.) is indicated at days 56 and 63 by an antibody symbol. Data are mean ± SEM, N=6-8 rats per group, *p<0.05 two-way ANOVA (**B**) Bar graph with scatter plot showing the area under the curve for each individual (n=8) from the indicated treatment groups. (**C**) Representative recordings of evoked action potentials recorded from rat small-diameter DRG neurons cultured from the indicated treatment groups in response to depolarizing current injection of 30, 60, and 90 pA. (**D**) Quantification of the number of current-evoked action potentials in response to 0-100 pA injected current. CRMP5 autoimmunity increased action potential firing compared to control DRG neurons. Anti-CD20 reversed this phenotype back to the level of control sensory neurons. *p<0.05, multiple Mann-Whitney tests. N=8-9 cells each, mean ± SEM.

## Discussion

Here, we report a previously unknown role of CV2/CRMP5-Abs in autoimmune painful neuropathy experienced by patients. These autoantibodies specifically target dorsal root ganglion (DRG) neurons and spinal cord laminae I/IIo, where nociceptive signals converge. Intrathecal or intra-paw application of sera containing these autoantibodies induces mechanical hypersensitivity in rats. At the cellular level, sensory neurons become hyperexcitable upon exposure to CV2/CRMP5-Abs. We identified epitopes on CRMP5 that are targeted by these autoantibodies, and successfully masked them to reduce DRG neuron sensitization and mechanical hypersensitivity. Additionally, the painful autoimmune neurological syndrome was replicated preclinically following DNA immunization. Notably, the anti-CD20 monoclonal antibody, named Rituximab in the clinic, emerged as a promising treatment option for patients with CRMP5 autoimmune painful neuropathy. Overall, the data strongly suggests that pain symptoms in anti-CV2/CRMP5 patients result from auto-immunoreactivity with DRG neurons, which can be effectively managed with anti-CD20.

This study stems from the remarkable clinical observation that approximately 80% of patients with CV2/CRMP5- Abs exhibit painful neuropathy (2). This prevalence of pain in a rare disease surpasses what is typically reported in common pain conditions, including diabetic neuropathy (42%) (6) and chemotherapy-induced neuropathy (60%)(8). Pain management for CV2/CRMP5-Abs patients necessitates at least two pain medications, with opioids being used in 39% of cases (2). Notably, evidence suggests that pain in anti-CRMP5 patients can be alleviated by reducing antibody levels (2). These findings strongly implicate the autoantibodies as direct contributors to heightened pain sensations in patients. To investigate this, we employed two approaches to directly assess whether CV2/CRMP5-Abs could enhance pain sensitivity: 1) we removed CV2/CRMP5-Abs from patient sera through adsorption on a purified CRMP5 substrate, and 2) we targeted the Fab regions of human autoantibodies using peptides that mask the epitopes on CRMP5. Both approaches effectively neutralized the functional impact of CV2/CRMP5-Abs on DRG neuron excitability and pain. Intriguingly, CV2/CRMP5-Abs induced mechanical allodynia, whether applied directly to the spinal cord or injected into the hindpaw. These findings lead us to conclude that CV2/CRMP5-Abs are pathogenic and that therapies aimed at reducing autoantibody levels in patients hold promise for mitigating their pain symptoms.

The auto-antigen target of CV2/CRMP5-Abs, CRMP5, is a cytosolic protein (11, 34). While these autoantibodies can recognize a cytoplasmic antigen in fixed DRG and spinal cord tissues (15), their ability to access the cytosolic CRMP5 protein under physiological conditions remains uncertain. Previous reports have demonstrated that autoantibodies can penetrate cells to bind to intracellular targets, even in the case of nuclear antigens (e.g., anti-Hu, Sm, or La) (35–37). In a separate study involving rats, peripherally injected rabbit IgG was transported into neurons within the spinal cord, further illustrating that antibodies can penetrate neuronal cytoplasm in vivo (38). A post-mortem study in patients with paraneoplastic syndrome revealed positive immunofluorescence for circulating autoantibodies in Purkinje and DRG neurons (39). Specifically concerning DRG neurons, studies have suggested that following immunization in rats, antibodies were sequestered and retained within their soma (40). These intriguing findings raise the possibility that sensory neurons are particularly sensitive to autoantibodies targeting neuronal antigens, emphasizing the intricate relationship between the immune and nervous systems. Based on our results, we hypothesize that the elevated prevalence of pain in autoimmune diseases, such as anti-CV2/CRMP5 autoimmune encephalitis, may be linked to the intracellular retention of autoantibodies in sensory neurons, ultimately contributing to pain sensitization. This hypothesis will require further exploration.

We demonstrated that CV2/CRMP5 Abs can bind their target in DRG sensory neurons to induce hyperexcitability and pain. Although CRMP5 has not previously been linked to pain, these data suggest that it may play a role in sensory neuron function. Another member of the CRMP family, CRMP2 has been extensively described to be phosphorylated and SUMOylated in chronic pain to increase the function of voltage gated ion channels NaV1.7 and CaV2.2 supporting hyperexcitability (18, 41–45). During neuronal polarization, CRMP5 is known to antagonize CRMP2 function promoting microtubule polymerization (13). In sensory neurons, CRMP5 could play a similar role against CRMP2 function by triggering ion channel internalization. Internalized channels are not able to function which would put the brakes on sensory neuron excitability. CV2/CRMP5 Abs would block the antagonistic action of CRMP5 thereby allowing for facilitated action potential firing in sensory neurons.

Our comprehensive investigations using calcium imaging and electrophysiology revealed that CV2/CRMP5-Abs exert a specific impact on sensory neuron function. Constellation pharmacology ruled out the possibility that heightened sensitivity to heat, cold, itch, or inflammation by DRG neurons could account for pain in patients with these autoantibodies. Instead, we found that DRG neurons exposed to CV2/CRMP5-Abs exhibited increased action potential firing. Interestingly, our findings diverged when examining the effect on rheobase, a measure of neuronal excitability. While human sera yielded conflicting results regarding rheobase alteration by CV2/CRMP5- Abs, our rat data consistently demonstrated that CRMP5 autoimmunity reduced the rheobase in both animals of both sexes. This suggests that CV2/CRMP5-Abs have a dual impact: lowering the threshold for firing and increasing the number of action potentials generated in response to stimuli. Notably, we correlated excitability with mechanical hypersensitivity throughout our experiments. Hyperexcitability, a hallmark of neuropathic pain observed in both rodent models and patients (20, 21), implies that pain symptoms in individuals with CV2/CRMP5-Abs may be more amenable to drugs aimed at reducing neuronal activity. In the absence of clinically available drugs aimed at silencing sensory neurons, our screening campaign revealed enhanced benefits from amitriptyline and duloxetine in reversing CRMP5 autoimmunity-induced mechanical hypersensitivity.

Limitations of our study include the restricted and variable availability of CV2/CRMP5-Abs positive sera in the biobank. Consequently, we were unable to systematically test all patients’ sera, which might have resolved the conflicting results observed in rheobase measurements. Additionally, it is uncertain if our collection might exhibit bias on the basis of patient sex though sex-based prevalence associated with the development of CV2/CRMP5- Abs have not been reported (2, 15). Despite these limitations, our epitope profiling revealed that all samples identified similar epitopes on CRMP5, suggesting equivalence from this perspective. We deliberately chose not to account for the titer of autoantibodies in our samples, as prior studies have not consistently correlated these measures with symptom severity in patients. Our strategies for depleting autoantibodies or blocking the Fab domain demonstrated that other serum components had limited impact on our pain and excitability studies, reinforcing the pathological role of CV2/CRMP5-Abs. Our findings with human sera align with data showing that depleting B-lymphocytes using the anti-CD20 monoclonal antibody Rituximab could serve as a long-term treatment for CRMP5 autoimmune painful neuropathy. Notably, patients with anti-CV2/CRMP5 autoimmunity are often diagnosed with cancer. Treatment with anti-CD20 bears the potential to synergize with classical chemotherapeutics (33) which could help with the elimination of primary tumors additionally to relieving autoimmune pain. Furthermore, pain is a significant comorbidity in cancer patients, and addressing pain in anti- CV2/CRMP5 patients can profoundly impact their quality of life and survival post-cancer diagnosis.

This first of its kind study directly assesses the function of an autoantibody targeting an intracellular protein. We demonstrate that CV2/CRMP5-Abs are drivers of hypersensitivity to mechanical stimuli and hyperexcitability in DRG neurons. We replicated CRMP5 autoimmunity in rats which provided a unique preclinical platform for investigating future treatment options to relieve pain in patients. Our work identifies clinically used painkillers with a superior ability to alleviate pain in patients with CV2/CRMP5-Abs. We propose a novel therapeutic approach by repurposing anti-CD20 monoclonal antibodies to treat pain long term in patients with CRMP5 autoimmunity.

## Acknowledgements

L.M. and H.J.S are co-first authors. Their contribution to this research was equal and are named in alphabetical order. We thank Mid America transplant and Dr. Grant Kolar for their support in facilitating access to human DRG. This research was supported by startup funds from Saint Louis university, National Institutes of Health NINDS R01NS119263, R01NS119263-04S1 to A.M. and NINDS R01NS098772 and R01NS120663, NIDA DA042852 to R.K. We are grateful to Genentech for sharing their anti-CD20 monoclonal antibody. Most authors completed this work while at the University of Arizona. LDD, VR and JH are supported by a public grant overseen by the *Agence Nationale de la Recherche* (ANR; French research agency) as part of the *Investissements d’Avenir* program (ANR-18-RHUS-0012), also performed within the framework of the LABEX CORTEX of the Université Claude Bernard Lyon 1 (program *Investissements d’Avenir*, ANR-11-LABX-0042, operated by the ANR), and also supported by the European Reference Network RITA. Current affiliations are as follows, HJS: Pittsburgh Center for Pain Research, University of Pittsburgh, Pittsburgh, PA 15213, USA; KG, SLL, ACR and RK : Department of Department of Pharmacology & Therapeutics, College of Medicine, University of Florida, Gainesville, FL 32610 USA; CT: The National and Local Joint Engineering Laboratory of Animal Peptide Drug Development, College of Life Sciences, Hunan Normal University, Changsha, 410081, China; DR: Department of Pharmacology, School of Pharmacy, Chongqing Medical University, Chongqing 400016, China; SSB : Medical Scientist Training Program (MSTP), Mayo Clinic, Rochester, MN, USA.

## Methods

### Patients

Sera used in this study were collected as part of the standard diagnostic procedures in the French reference center on paraneoplastic neurological diseases and then stored for research purposes as part of the Neurobiotec biobank in Lyon, France. De-identified samples with informed consent of the patients were transferred to the laboratory overnight via Fedex. Research using human sera without patient personal information was performed under an Institutional review board exemption 4 at the University of Arizona and at Saint Louis University.

### Animals

Adult Sprague-Dawley rats (Pathogen-free male and female 100–250 g, Envigo, Placentia, CA) were kept in light (12-h light: 12-h dark cycle; lights on at 07:00 h) and temperature (23 ± 3°C) controlled rooms. Standard rodent chow and water were available *ad libitum.* All animal use was conducted in accordance with the National Institutes of Health guidelines, and the study was conducted in strict accordance with recommendations in the Guide for the Care and Use of Laboratory Animals of Saint Louis University (Protocol #: 3014). All behavioral experiments were performed by experimenters who were blinded to treatment groups.

### Human Dorsal Root ganglia

Human DRG were obtained from Mid America Transplant at St. Louis University within an hour of cross-clamp and immediately picked up and processed by laboratory personnel. All studies involving human tissues have been classified for an IRB exemption #4 at Saint Louis University. DRGs from one female donor were fixed by immersion in 10% formalin for at least one day. After fixation, the DRGs were trimmed of their connective tissue and roots, cut in half, prior to embedding into paraffin blocks. The paraffin blocks were sectioned in 5 µm slices. Deparaffination is carried out by three xylene washes for 5 minutes each, followed by two 100% alcohol washes for 1 minute each, then two 95% alcohol washes for 1 minute each, and finally by washing in running water for 2 minutes. For antibody staining, the slides were washed twice with phosphate buffered saline (PBS) for 5 minutes each at room temperature then exposed for 30 minutes to a blocking solution containing 5% donkey serum and 1% BSA in PBS at room temperature. The primary antibody against CRMP5 (1:100, from (17)) was then incubated in blocking buffer, overnight at 4°C in a humidified chamber. The slides were then washed 3 times, 15 minutes with PBS at room temperature before adding the secondary antibody solution (donkey anti-rabbit, alexa488, 1/300) for one hour at room temperature. Slides were then washed with PBS 3 times for 15 minutes at room temperature. Finally, slides were mounted in ProLong Gold (Cat# P36930, Thermo Fisher Scientific) to protect from fading and photobleaching. To evaluate background due to the secondary antibody alone, primary antibody was omitted. Immunofluorescent micrographs were acquired on a Leica SP8 inverted microscope using a 10x dry objective. The freeware image analysis program Image J (http://rsb.info.nih.gov/ij/) was used for extracting representative pictures after contrast enhancement for better visualization.

### Materials and Reagents

All peptides (98% purity) were purchased from Genscript (Piscataway, NJ). All chemicals, unless noted were purchased from Sigma (St. Louis, MO). Fura-2 AM was obtained from Life technologies. Subcellular proteome extraction kit (Cat# 539790, Millipore). Antibodies used are as follows: CRMP5 from (17), Goat anti-Human IgG Cross-Adsorbed, Alexa Fluor™ 488 (Cat# A11013, Life technologies), Goat anti-Rabbit IgG, Alexa Fluor™ 555 (Cat# A21429, Life technologies), Rabbit anti-Human IgG DyLight™ 800 (Cat# SA5-10116, Thermofisher). For the appraisal of common painkillers drugs were as follows: ibuprofen (100 mg/kg, Cat# I4883, Sigma), acetaminophen (200 mg/kg, Cat# A3035, Sigma), gabapentin (100 mg/kg, Cat# G154, Sigma), amitriptyline (50 mg/kg, Cat# A8404, Sigma), duloxetine (30 mg/kg, Cat# SML0474, Sigma) and morphine (10 mg/kg, Cat# M8777, Sigma). The selected medication doses were determined from previously documented animal studies, and we opted for oral administration as it is the predominant method utilized in clinical settings (46–50).

### Purification of CRMP5-GST

CRMP5-GST fusion protein was purified similarly to a previously described protocol (Francois-Moutal et al., 2015). BL21 *Escherichia coli* cells (New England Biolabs, Ipswich, MA) expressing recombinant CRMP5-GST were resuspended in 50 mM HEPES, pH 7.5, 500 mM NaCl, 10% glycerol (vol./vol/), 0.5 mM Tris(2- carboxyethyl)phosphine hydrochloride (TCEP), supplemented with complete EDTA-free protease inhibitors (Roche, Basel, Switzerland). Disruption of the bacteria was performed by sonication, and the lysate was centrifuged at 4°C for 45 min at 20,000×g. The supernatant was loaded on a GST-Trap HP column (Cytiva, Marlborough, MA) equilibrated with 50 mM HEPES pH 7.5, 500 mM NaCl, 10% glycerol, 0.5 mM TCEP. After a washing step with 50 mM HEPES, pH 7.5, 5 mM Glutathione, 500 mM NaCl, 10% glycerol, 0.5 mM TCEP, CRMP5-GSTwas eluted with a step gradient up to 40mM Glutathione. The eluted protein was concentrated with Amicon Ultra 15 centrifugal filters (Regenerated cellulose 10 000 NMWL; Merck Millipore, Darmstadt, Germany), aliquoted and flash-frozen on dry ice and stored at −80°C until use. Protein concentration was determined by a Pierce assay using BSA as a standard. The purity of the protein was verified with SDS-PAGE.

### Immunohistochemistry

Freshly prepared adult rat spinal cord was fixed in 4% paraformaldehyde (EM-15713-S, Euromedex, Sousselweyrsheim, France) for 1h, frozen, and sliced into 10 µm-thick sections. Tissue was permeabilized and blocked during 1h at RT in a blocking buffer containing PBS supplemented with 5% (v/v) normal goat serum, 2% (m/v) bovine serum albumin and 0.25% (v/v) Triton X-100. Double immunolabeling was performed using patient serum (1:100) and rabbit anti-CRMP5 antibody (1:400, (17)) in blocking buffer during 2h at RT. Tissue was washed 3 times 10 min, at RT with 1ml of PBS, and revealed with appropriate Alexa fluorophore-conjugated secondary antibodies (1:1000 in blocking buffer, A11013 and A21429, Thermofisher, Courtaboeuf, France) and DAPI (10 µg/mL) during 1h at RT. Photos of immunolabeling were acquired on Axio Scan.Z1 (Carl Zeiss SAS, Rueil Malmaison, France) with 20x, 0.8 NA, Plan-Apochromat objective (Carl Zeiss SAS, Rueil Malmaison, France). Images were visualized and analyzed by means of ZEN (Blue) software (Carl Zeiss SAS, Rueil Malmaison, France) and ImageJ.

### Proximity Ligation assay

PLA was performed to visualize the close proximity (less than 30 nm) between the human anti-CV2 autoantibodies and their target CRMP5 in cultured DRG neurons. DRG neurons were incubated overnight with the indicated sera diluted 1:100 before fixation using 4% paraformaldehyde for 20 min at RT. Blocking and permeabilization was done by incubating the cells with PBS, 0.1 % triton X-100 with 3 % BSA for 30 minutes at RT. Fixed DRG neurons were incubated with anti-CRMP5 antibody for 1 hour at RT in PBS, 0.1 % triton X-100, 3 % BSA before 3 washes in PBS, 0.1 % triton for 5 min at RT. The proximity ligation reaction and visualization of signal was performed according to the manufacturer’s protocol using the Duolink Detection Kit with PLA PLUS and MINUS probes for rabbit and human antibodies (Sigma). DAPI stain was used to detect cell nuclei. Immunofluorescent micrographs were acquired on a Nikon Eclipse Ti/U microscope with a photometrics cooled CCD camera CoolSNAP ES2 (Roper Scientific, Planegg, Germany) controlled by NIS Elements software (version 4.20, Nikon instruments), using a 60X plan Apo 1.40 numerical aperture objective.

### Enzyme-linked immunosorbent assay-based CRMP5-anti-CV2 binding assay

96-well plates (Nunc MaxiSorp, Thermo Scientific) were coated with CRMP5-GST (200 ng per well) and incubated at room temperature overnight. The next day, the plates were washed and blocked with 3% BSA to minimize nonspecific adsorptive binding to the plates. The plates were incubated at room temperature with serum from patients (diluted 1:100 in PBS) with the indicated peptide at 100 ng/ml and left on gentle shaking for 2 hours. The plates were then washed with PBS containing 0.5% Tween-20. The bound anti-CV2 was detected by HRP-conjugated secondary antibody. Tetramethylbenzidine (R&D Systems) was used as the colorimetric substrate. The optical density of each well was determined immediately, using a microplate reader (Multiskan Ascent, Thermo) set to 450 nm with a correction wavelength of 570 nm.

### Peptide synthesis and spotting on a peptide array

Standard 9-fluorenylmethoxy carbonyl (Fmoc) chemistry was used to synthesize the peptides and spot them onto Celluspots nitrocellulose disks prederivatized with a polyethylene glycerol spacer (Intavis). Peptide synthesis was done using the Respep peptide synthesizer using established protocols from Intavis. After synthesis, a side chain deprotection step was done in 80% trifluoroacetic acid (TFA), 3% triisopropylsilane (TIPS), 12% dichloromethane (DCM), 5% H2O for 2 hours at room temperature. Next, the celluspots were solubilized in 88.5% TFA, 4% trifluoromethanesulfonic acid (TFMSA), 2.5% TIPS, 5% H2O overnight at room temperature before precipitation by adding 4:1 Ice cold tert-butyl-methyl ether. Cellulose-peptide conjugates were pelleted by a 5000xg, 10 min, 0°C centrifugation step. After removing the supernatant, the pellets were allowed to dry until achieving a gel-like texture at room temperature and resuspended in 100% DMSO. The peptides were spotted on 20 membranes fitted on microscope glass slides (Intavis) using an Intavis MultiPep robot. All chemicals were HPLC grade and obtained from Sigma.

### Hybridization and immunoblotting of peptide arrays

Celluspots slides were washed in TBST (50 mM Tris-HCl, pH 7.4, 150 mM NaCl, 0.1 % Tween 20) for 10 min, 5% (mass/vol) non-fat dry milk and then blocked for 1 h at room temperature (RT) with gentle shaking in TBST containing 5% (w/v) nonfat dry milk. Sera were diluted 100 times in TBST, 5% nonfat dry milk and incubated on the slide overnight. Peptide arrays were washed 3 times for 5 min at RT with TBST and incubated with the antibody Rabbit anti-Human IgG DyLight™ 800 (Cat# SA5-10116, Thermofisher, 1/1000) diluted in TBST, 5% BSA for 2h at RT. The arrays were washed 3 times, 5 min in TBST, and visualized by infrared fluorescence. All arrays contained quadruplicates that were averaged and then normalized on the maximum signal on the array for each patient.

### Acute dissociation and culture of dorsal root ganglia (DRG) neurons

Dorsal root ganglia (DRG) were dissected from Sprague-Dawley rats. In brief, removing dorsal skin and muscle and cutting the vertebral processes parallel to the dissection stage exposed DRGs. DRGs were then collected, trimmed at their roots, and digested in 3 ml bicarbonate free, serum free, sterile DMEM (Cat# 11965, Thermo Fisher Scientific, Waltham, MA) solution containing neutral protease (3.125 mg/ml, Cat#LS02104, Worthington, Lakewood, NJ) and collagenase Type I (5 mg/ml, Cat# LS004194, Worthington, Lakewood, NJ) and incubated for 45 min at 37°C under gentile agitation. Dissociated DRG neurons (∼1.5 x 10^6^) were then gently centrifuged to collect cells and washed with DRG media DMEM containing 1% penicillin/streptomycin sulfate from 10,000 µg/ml stock, 30 ng/ml nerve growth factor, and 10% fetal bovine serum (Hyclone). Cells were plated onto poly-D-lysine - and laminin-coated glass 12- or 15-mm coverslips. For rat DRG culture small cells were considered to be ∼ < 30 µm diameter in size. All cultures were used within 48 hours.

### Calcium imaging

DRG neurons were loaded at 37°C with 3 μM Fura-2AM (Cat#F-1221, Life technologies, stock solution prepared at 1 mM in DMSO, 0.02% pluronic acid, Cat#P-3000MP, Life technologies) for 30 minutes (Kd= 25 μM, λex 340, 380 nm/λemi 512 nm) to follow changes in intracellular calcium ([Ca^2+^]c) in Tyrode’s solution (at ∼310 mOsm) containing 119 mM NaCl, 2.5 mM KCl, 2 mM MgCl2, 2 mM CaCl2, 25 mM HEPES, pH 7.4 and 30 mM glucose. All calcium-imaging experiments were done at room temperature (∼23°C). Baseline was acquired for 1 minute followed by stimulation (15 sec) with an excitatory solution (at ∼310 mOsm) comprised of 32 mM NaCl, 90 mM KCl, 2 mM MgCl2, 2 mM CaCl2, 25 mM HEPES, pH 7.4 and 30 mM glucose. Fluorescence imaging was performed with an inverted microscope, Nikon Eclipse T*i*-U (Nikon Instruments Inc.), using a Nikon Super Fluor MTB FLUOR 10x 0.50 NA objective and a Photometrics cooled CCD camera CoolSNAP ES^2^ (Roper Scientific) controlled by NIS Elements software (version 4.20, Nikon instruments). The excitation light was delivered by a Lambda-LS system (Sutter Instruments). The excitation filters (340±5 nm and 380±7 nm) were controlled by a Lambda 10-2 optical filter change (Sutter Instruments). Fluorescence was recorded through a 505 nm dichroic mirror at 535±25 nm. To minimize photobleaching and phototoxicity, the images were taken every 10 seconds during the time-course of the experiment using the minimal exposure time that provided acceptable image quality. The changes in [Ca^2+^]c were monitored by following the ratio of F340/F380, calculated after subtracting the background from both channels.

### Constellation pharmacology

DRG neurons were loaded at 37°C with 3 μM Fura-2AM for 30 minutes in Tyrode’s solution. After a 1-minute baseline measurement Ca^2+^ influx was stimulated by the addition of the following receptor agonists: 400 nM menthol, 50 µM histamine, 10 µM adenosine triphosphate (ATP), 200 µM allyl isothiocyanate (AITC), 1 mM acetylcholine (Ach), 100 nM capsaicin diluted in Tyrode’s solution. At the end of the constellation pharmacology protocol, cell viability was assessed by depolarization-induced Ca^2+^ influx using and an excitatory KCl solution comprised of 32 mM NaCl, 90 mM KCl, 2 mM MgCl2, 2 mM CaCl2, 25 mM HEPES, pH 7.4, 30 mM glucose. After the 1-minute baseline measurement, each trigger was applied for 15-seconds in the order indicated above in 6- minute intervals. Following each trigger, bath solution was continuously perfused over the cells to wash off excess of the trigger. Fluorescence imaging was performed under the same conditions noted above for calcium imaging. A cell was defined as a ‘responder’ if its fluorescence ratio of 340nm/380nm was greater than 10% of the baseline value calculated using the average fluorescence in the 30 seconds preceding application of the trigger.

### Measurement of action potentials using whole-cell current-clamp electrophysiology

Patch-clamp recordings were performed at room temperature (22–24°C). For current-clamp recordings the external solution contained (in millimolar): 154 NaCl, 5.6 KCl, 2 CaCl2, 1 MgCl2, 10 D-Glucose, and 8 HEPES (pH 7.4 adjusted with NaOH, and mOsm/L= 300). The internal solution was composed of (in millimolar): 137 KCl, 10 NaCl, 1 MgCl2, 1 EGTA, and 10 HEPES (pH 7.3 adjusted with KOH, and mOsm/L= 277). At room temperature (22–24°C), a tight seal with the cell membrane was established by applying negative pressure to obtain a seal resistance > 1GΩ. A brief pulse of negative pressure was then applied to rupture the cell membrane and establish the whole-cell patch clamp configuration. Neurons were initially held at -60 mV in voltage clamp mode to measure seal quality before switching to current clamp mode where the current injection was immediately set to 0 pA to measure the resting membrane potential. DRG neurons with a resting membrane potential (RMP) more hyperpolarized than −40 mV, stable baseline recordings, and evoked spikes that overshot 0 mV were used for experiments and analysis. The action potentials were evoked by current injection steps from 0–120 pA with an increment of 10 pA in 300 ms. Rheobase was measured by injecting currents from 0 pA with an increment of 10 pA in 50 ms. Analyses were performed by using Fitmaster software (HEKA) and Origin 9.0 software (OriginLab).

Pipettes were pulled from standard wall borosilicate glass capillaries (Sutter Instruments) with a horizontal puller (Model P-97, Sutter Instruments). The resistance of the pipettes when filled with internal solution and immersed in the recording bath ranged from 2 to 4 MΩ. Recordings were performed from small DRG neurons with capacitance between 10 and 35 pF (∼18-33 μm). Series resistance under 7 MΩ was deemed acceptable. All experiments had a series resistance compensation between 60-90 %. Signals were filtered at 10 kHz and digitized at 10–20 kHz. Analyzes were performed by using Fitmaster software (HEKA) and Origin 9.0 software (OriginLab).

### Indwelling intrathecal catheter

Rats were anesthetized (ketamine/xylazine anesthesia, 80/12 mg/kg i.p., Sigma) and placed in a stereotaxic apparatus. The cisterna magna was exposed and incised, and an 8 cm catheter (PE-10, Stoelting) was implanted as previously reported, terminating in the lumbar region of the spinal cord (51). Catheters were sutured (3-0 silk suture) into the deep muscle and externalized at the back of the neck; skin was closed with autoclips and behavioral evaluations were performed after a 5-7 day recovery period. Rats were injected with the indicated serum of their IgG depleted counterparts. IgG depletion was achieved using protein G dynabeads (Cat# 10004D, ThermoFisher) incubated with the whole serum for 2 hours at 4°C. Protein G dynabeads captured all the igG in the serum and the supernatant was considered depleted of IgGs.

### Testing of allodynia

The assessment of tactile allodynia (i.e., a decreased threshold to paw withdrawal after probing with normally innocuous mechanical stimuli) consisted of testing the withdrawal threshold of the paw in response to probing with a series of calibrated fine (von Frey) filaments. Each filament was applied perpendicularly to the plantar surface of the paw of rats held in suspended wire mesh cages. The withdrawal threshold was determined by sequentially increasing and decreasing the stimulus strength (the “up and down” method), and data were analyzed with the nonparametric method of Dixon, as described by Chaplan et al (52) and expressed as the mean withdrawal threshold.

### Measurement of thermal withdrawal latency

The method of Hargreaves et al. (53) was used to evaluate thermal sensitivity. Rats were acclimated within Plexiglas enclosures on a clear glass plate maintained at 23°C. A radiant heat source (high-intensity projector lamp) was focused onto the plantar surface of the hind paw. When the paw was withdrawn, a motion detector halted the stimulus and a timer. A maximal cutoff of 33.5 sec was used to prevent tissue damage.

### Mechanical conflict-avoidance (MCA) assay

Voluntary mechanical conflict-avoidance was measured using the Coy mechanical conflict-avoidance system (30). The MCA apparatus is made of a dark (safe) chamber, a sharp probe field and a bright (∼4800 lux, aversive/unsafe) chamber. The bright light serves as an aversive stimulus signaling an unsafe environment for rats. Rats have to cross a field of sharp probes (height 3mm) to reach the safe dark area. Rats are placed in the bright compartment with the light turned off and the escape door closed. Following 15-s of dark acclimation, the compartment light is turned on for the duration of the test. The escape door is opened 20-s thereafter. Time to cross the probe bed is recorded using a stopwatch starting from the time the escape door opens. The cut-off time for this experiment was 3 min.

### Genetic immunization of rats to induce CRMP5 autoimmunity

We chose to use DNA immunization (29, 54) as a mean to induce an immune response via both T- and B-lymphocytes (54–57). Another benefit of this approach is that the methylated CpG rich domains carried by the plasmid act as adjuvants to activate dendritic cells leading to local production of the cytokines including interleukin (IL)-6, IL-12 and tumor necrosis factor (TNF)-α and increase immune cell recruitment at the immunization site (58). Rats received a unilateral intramuscular in the spinodeltoidus muscle 50 µg of plasmid DNA (either empty pCMV2-Flag or pCMV2-CRMP5-Flag (34) in 200 µl of saline) (59). This was followed by two booster injections administered two weeks (booster 1) and four weeks (booster 2) after the first injection.

### Ex vivo calcitonin gene–related peptide (CGRP) release from lumbar spinal cord

Rats were deeply anesthetized with 5% isoflurane and then decapitated. Two vertebral incisions (cervical and lumbar) were made to expose the spinal cord. Pressure was applied to a saline-filled syringe inserted into the lumbar vertebral foramen, and the spinal cord was extracted. Only the lumbar region of the spinal cord was used for the CGRP release assay. Baseline treatments (#1 and #2) involved bathing the spinal cord in Tyrode’s solution. The excitatory solution consisting of 90 mM KCl (#3) was used to evoke CGRP release from the spinal primary afferents. These fractions (10 minutes, 400 µL each) were collected for measurement of CGRP release. Samples were immediately flash frozen and stored in a -20 °C freezer. The concentration of CGRP released into the buffer was measured by enzyme-linked immunosorbent assay (Cat# 589001, Cayman Chemical, Ann Arbor, MI).

### Fluorescence-activated cell sorting to detect B-lymphocytes in rat blood

Blood was collected from the tail vein of rats and red blood cells were lysed in 158 mM NH4Cl for 5 minutes. Lysis was quenched with PBS prior to centrifugation at 310xg to pellet intact cells. This process was repeated once and then cells were resuspended in the antibody solution containing APC anti-rat CD45RA (Cat# 202313, Biolegend) and FITC anti-rat CD3 (Cat# 201403, Biolegend) diluted 1:100 in PBS with 2% FBS for 30 minutes at room temperature. Samples were washed 3 times with PBS for 5 minutes and spun at 310xg in between washes. Controls were made by omission of either or both antibodies. Cells were analyzed by the Flow Cytometry Research Core Facility at Saint Louis University.

### Rat Cytokine array

Serum from 6 rats was pooled and hybridized with the Proteome Profiler Rat XL Cytokine Array (Cat # ARY030, R&D systems) according to manufacturers instructions. Signal was captured on photographic films, scanned, and quantified with Un-Scan IT 7.0 (Silk scientific). All arrays were processed in parallel and imaged simultaneously. Signal was background subtracted and normalized to positive reference points. Cytokine profiling was calculated as a log2(fold change) compared to rats with CRMP5 autoimmunity.

### Statistical methods and data analysis

Graphing and statistical analysis was undertaken with GraphPad Prism (Version 9). All data sets were checked for normality using D’Agostino & Pearson test. Details of statistical tests, significance and sample sizes are reported in the appropriate figure legends and in **Table S1**. All data plotted represent mean ± SEM. For electrophysiological recordings: data was compared using Mann-Whitney tests (rheobase), One-way ANOVA with the Tukey post hoc test and Kruskal–Wallis test with Dunnett’s post hoc comparisons; for sensory neuron excitability, statistical differences between groups were determined using multiple Mann-Whitney test and Mann-Whitney test. Statistical significance of hypersensitivity was compared by Kruskal-Wallis test followed by the Dunn post hoc test. Behavioral data with a time course were analyzed by two-way ANOVA.

---

**Supplementary Figure 1:**
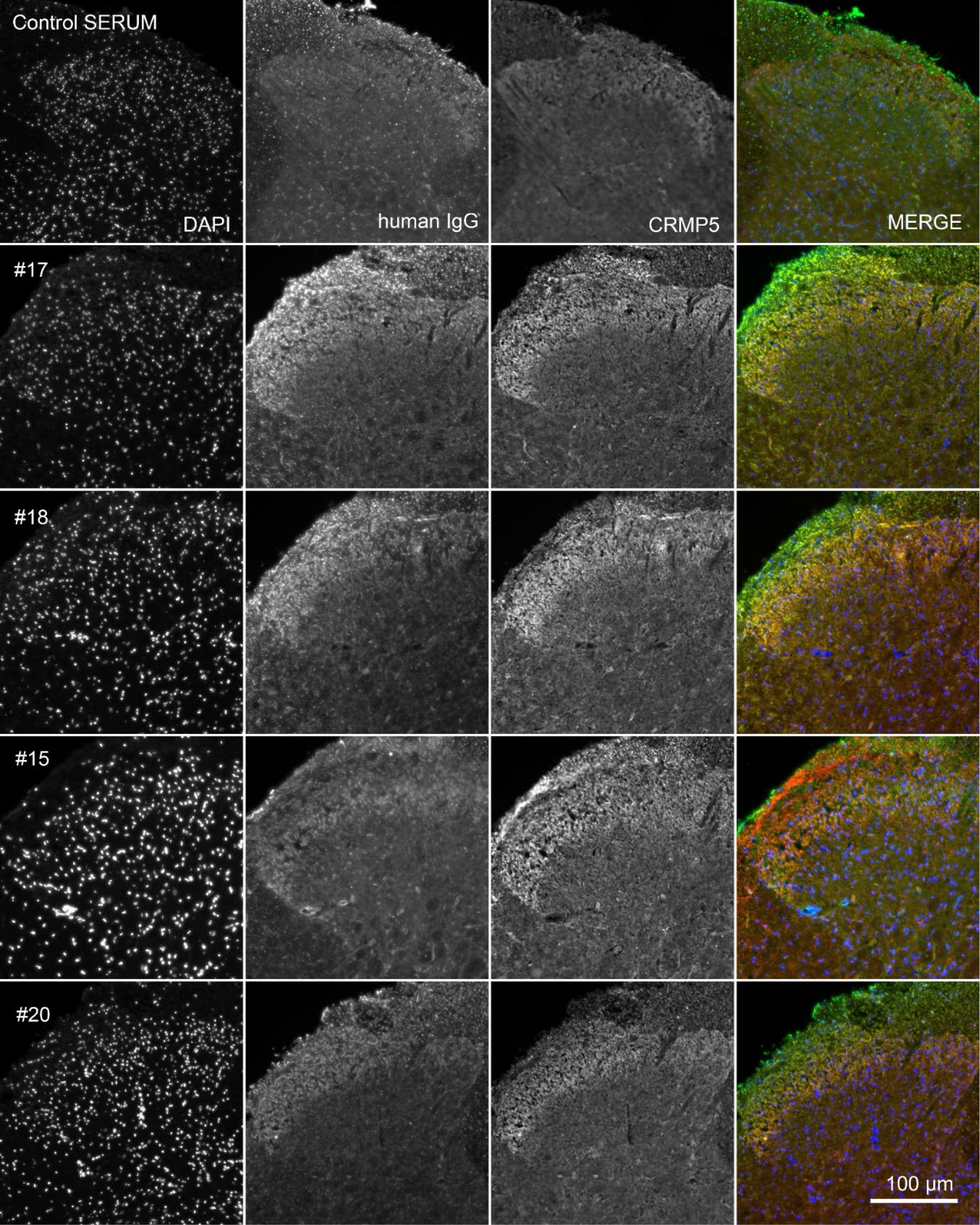
Anti-CV2 immunoreactivity colocalizes with CRMP5 in lumbar spinal cord. Micrographs of rat lumbar spinal cord co-immunolabelled with the indicated anti-CV2 or control sera and with an antibody against CRMP5. Anti-CV2 autoantibodies and anti-CRMP5 positively labelled and colocalized in superficial laminae in the dorsal horn of the lumbar spinal cord.

**Supplementary Figure 2:**
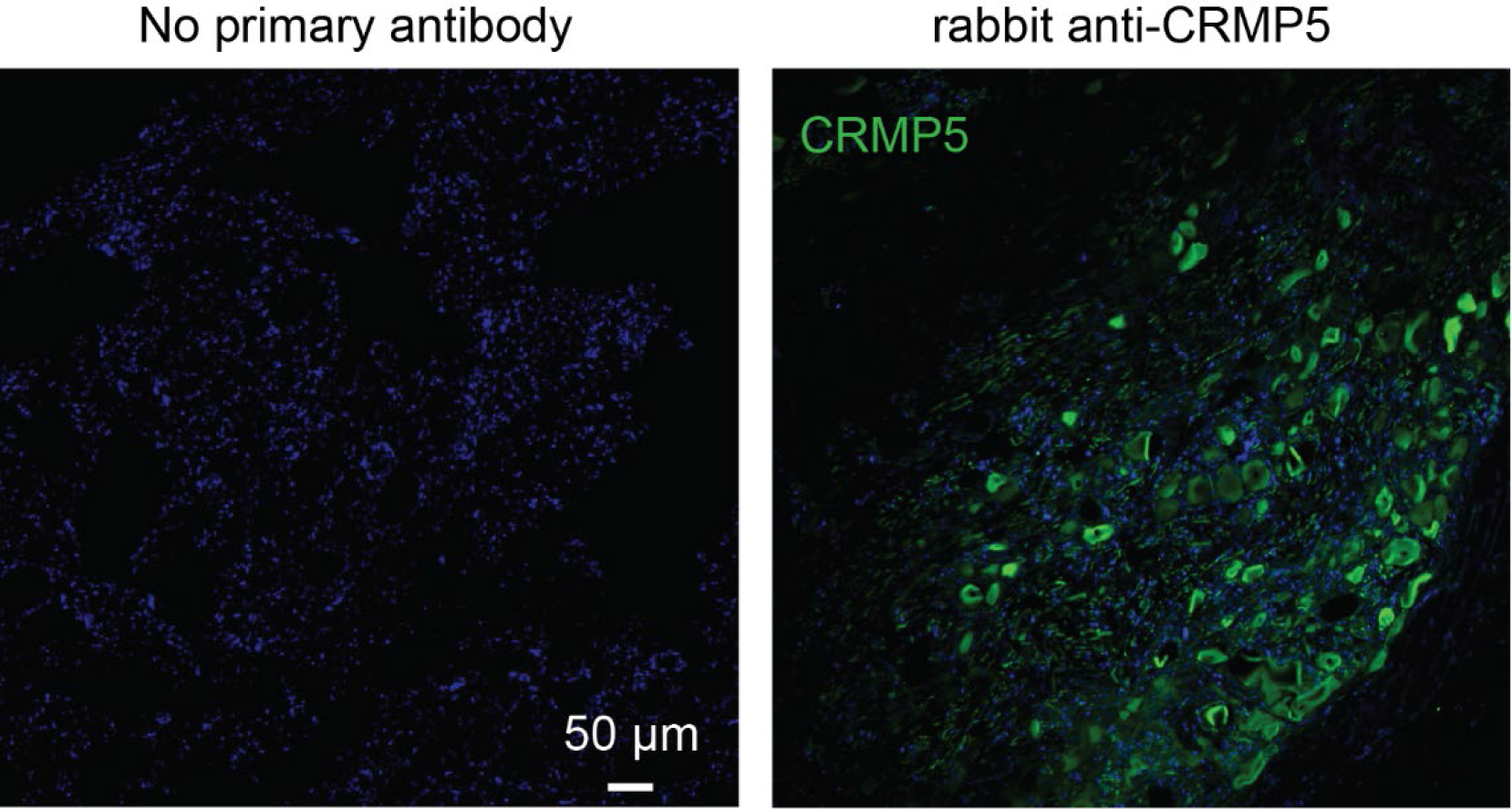
CRMP5 expression in sensory neurons in human DRG tissues. Human DRG were immunolabelled for CRMP5 (polyclonal from (16)). The left panel shows the negative control where primary antibodies were omitted. These data show that the target of autoantibodies CRMP5 is expressed in human DRG neurons. Nuclei were counterstained with DAPI. Scale bars: 50µm.

**Supplementary Figure 3:**
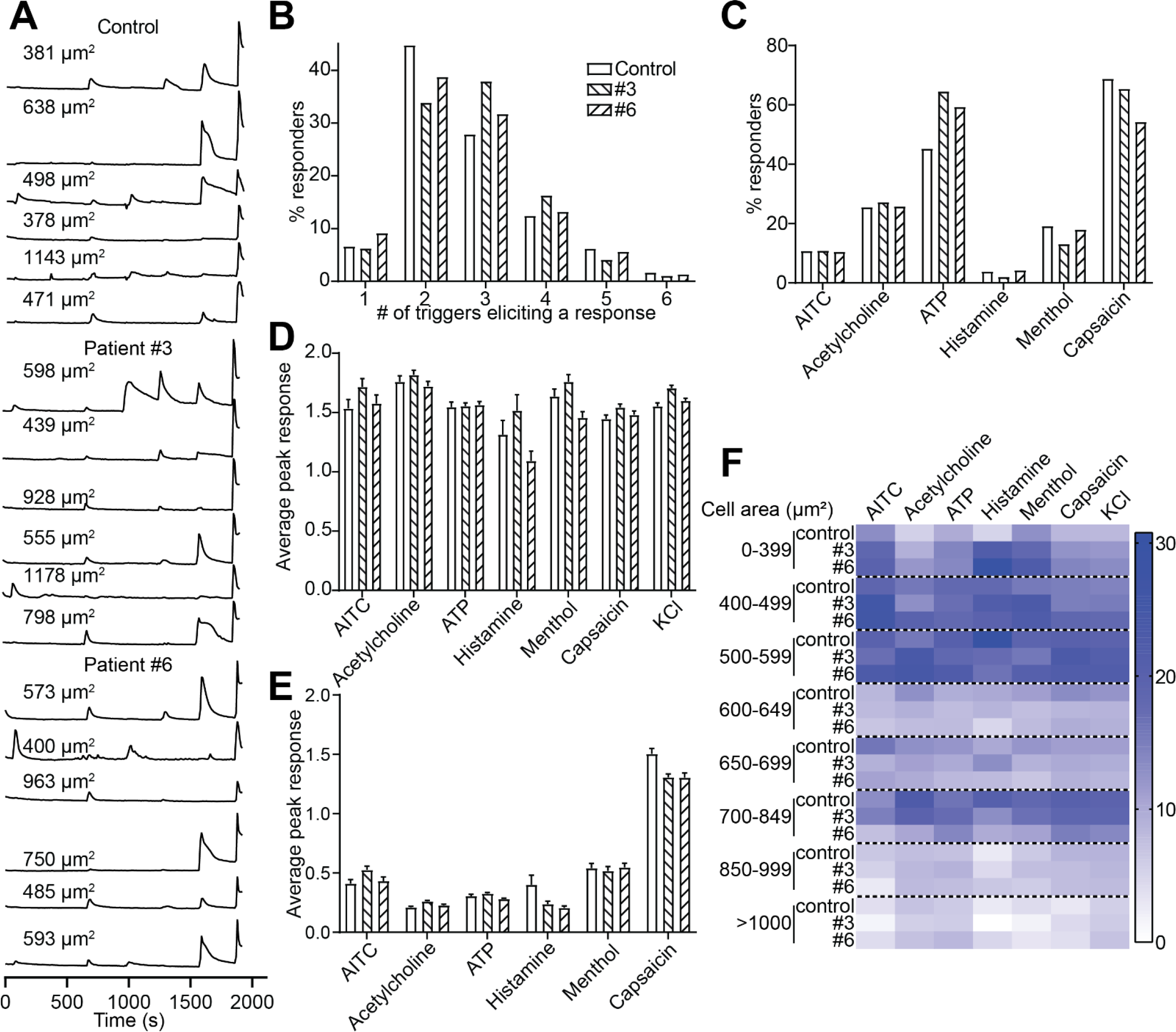
Functional “fingerprinting” of dorsal root ganglion (DRG) neuronal subclasses after exposure to anti-CV2 sera. (**A**) Representative traces of sensory neurons treated with the indicated sera (1/100 dilution, overnight) responding to constellation pharmacology triggers (menthol [400 nM], histamine [50 µM], ATP [10µM], AITC [200 µM], ACh [1 mM], capsaicin [100 nM], and KCl [90 mM]). Each trace represents an individual neuron; a typical experimental trial records the responses of .200 neurons concurrently. The x-axis represents time in seconds and the y-axis shows the relative intracellular calcium [Ca^2+^] in each DRG neuron (i.e., the F340/F380 ratio). (**B**) Percentage of DRG sensory neurons that responded to the indicated number of triggers. “1” indicates the neurons that only responded to none other than KCl stimulus. (**C**) Percentage of sensory neurons responding to major classes of constellation triggers. (**D**) Average peak responses are shown for calcium responses in sensory neurons after the indicated treatment, after stimulation by major classes of constellation triggers. (**E**) Average peak KCl-evoked response of sensory neurons after indicated treatment. (**F**) Heatmap of the size distribution of neurons responsive to each receptor agonist. In all analyses, the functional landscape of DRG neurons was not impacted by anti-CV2 exposure. Abbreviations for constellation triggers are as follows: AITC, allyl isothiocyanate; ATP, adenosine triphosphate; KCl, potassium chloride. Error bars indicate mean ± SEM. Data were acquired from a total of 3 independent experiments with an n=965 for the control serum and n=1748 (Patient #3) and n=1368 (Patient #6).

**Supplementary Figure 4:**
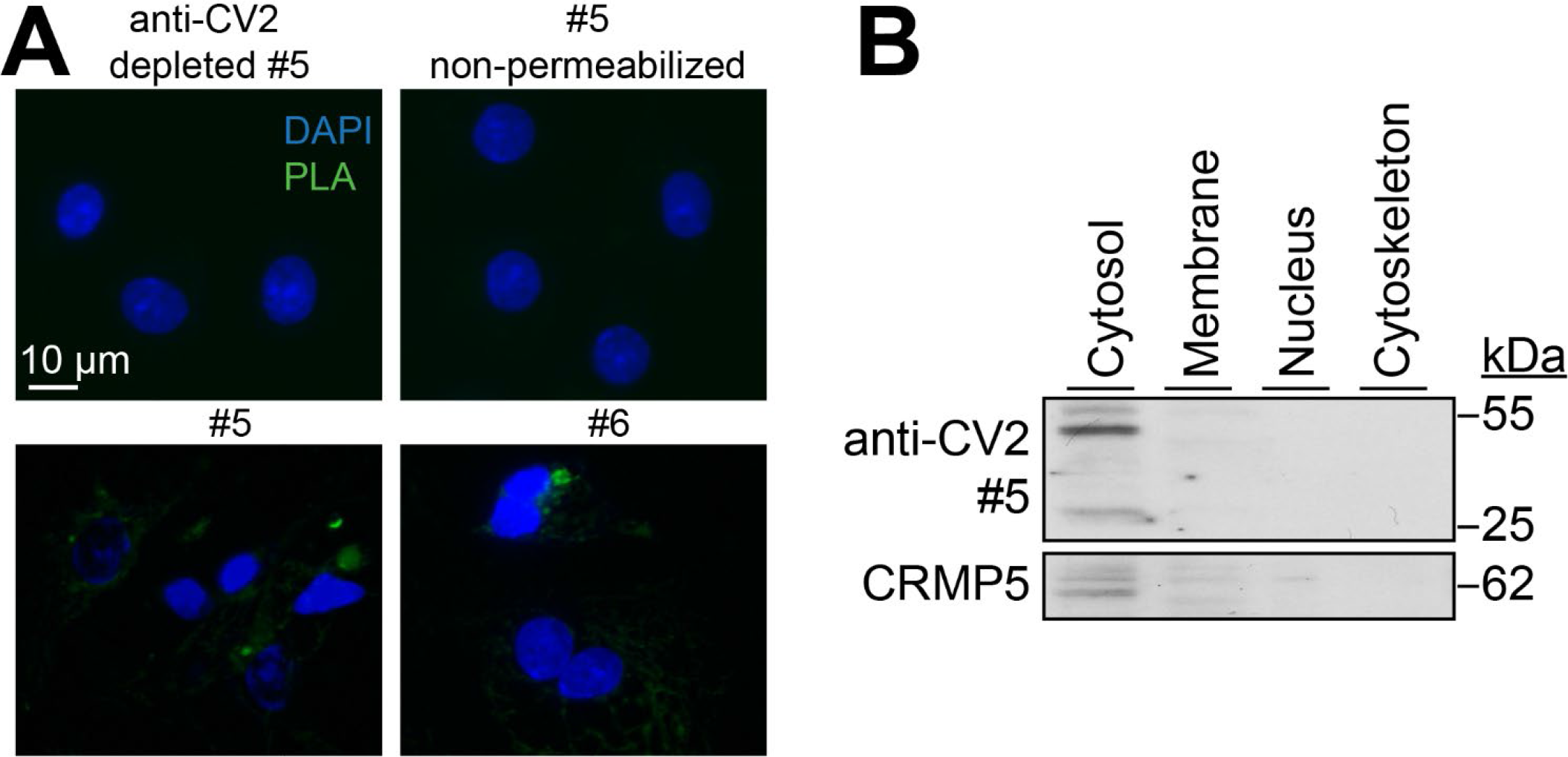
Anti-CV2 autoantibodies can enter sensory neurons to bind cytoplasmic CRMP5. (**A**) Representative micrographs showing the proximity ligation assay (PLA) signal of cultured adult rat DRG neurons incubated with anti-CV2 positive serum, co-stained for human IgG and CRMP5. Controls included anti-CV2 depleted serum from the same patient as well as non-permeabilized cells. The PLA signal (green) shows intracellular binding of human anti-CV2 antibody to cytosolic CRMP5. (**B**) Representative immunoblot of adult rat DRG neurons incubated with patient serum and fractionated to isolate the cytosolic, membrane, nuclear and cytoskeletal fractions. A positive signal was found for human IgG (heavy chain at ∼55 kDa, light chain at 25kDa) in the cytosolic fraction and weakly in the membrane fraction.

**Supplementary Figure 5:**
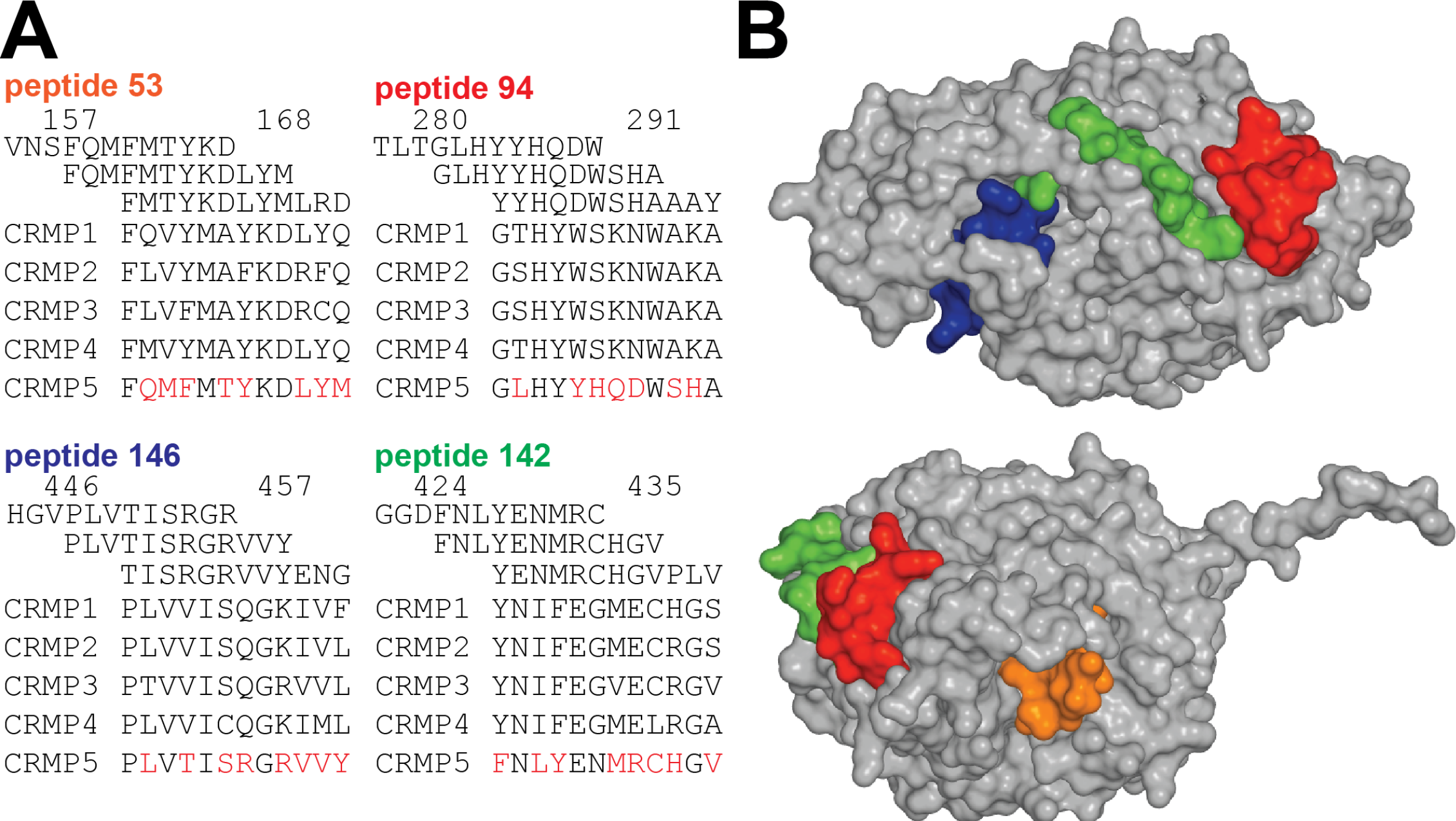
Anti-CV2 epitopes are unique and available on the surface of CRMP5. (**A**) Spatial representation of the epitopes identified in Figure 2A, localized at the surface of CRMP5, model based on PBDID:4B90 (25). (**B**) Alignment of the main epitopes found in **A** compared to the other members of the CRMP family of proteins. Epitopes are unique to CRMP5.

**Supplementary Figure 6:**
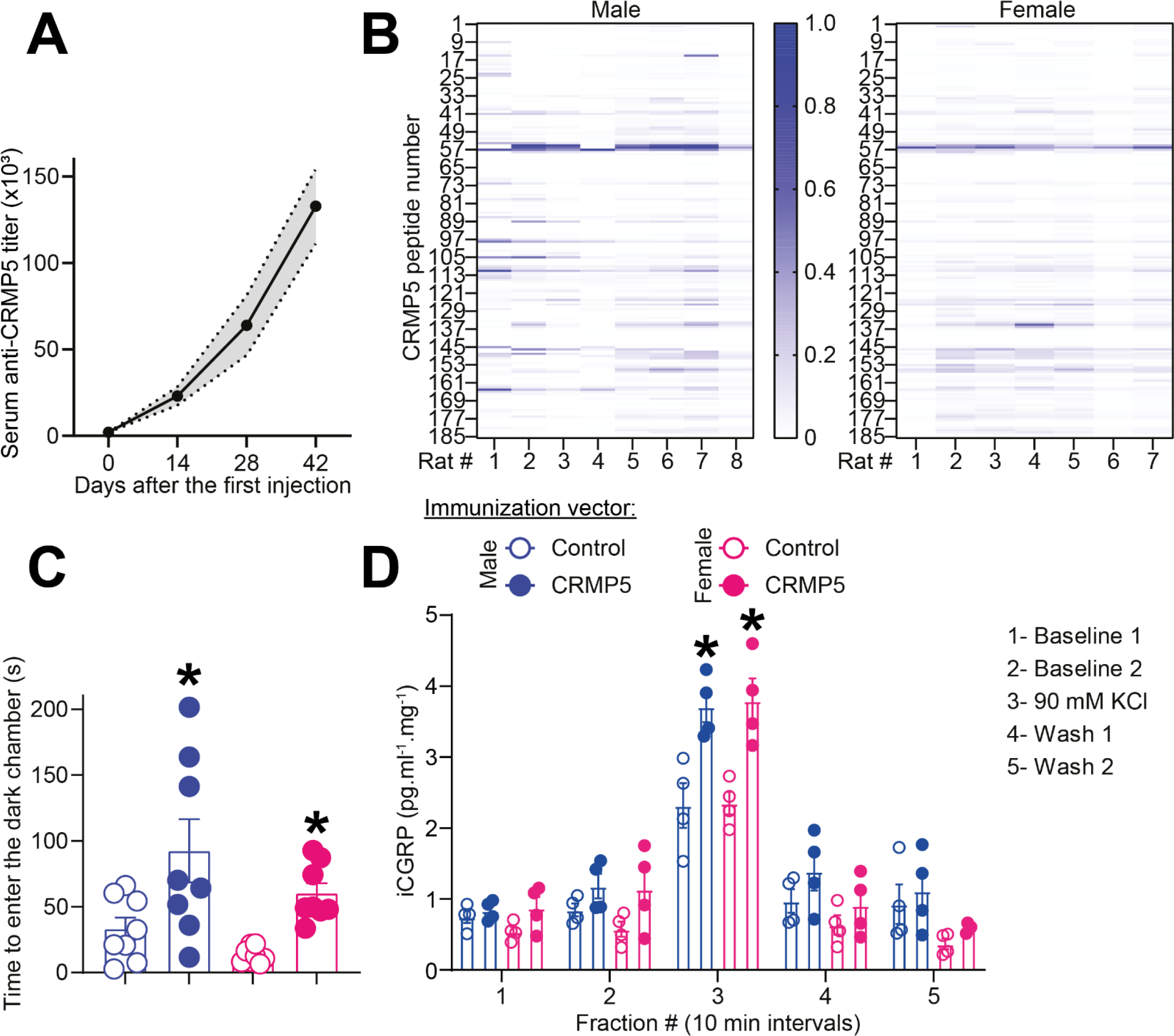
DNA immunization induces CRMP5 autoantibodies, mechanical pain, and increased neurotransmitter release from the spinal cord. (**A**) Graph showing anti-CRMP5 titers in rat serum measured by ELISA against purified CRMP5 protein. Anti-CRMP5 autoantibodies were found only in rats injected with the CRMP5 expressing plasmid. Serum titers increased after each booster injection. *p<0.05 compared to Control plasmid injected rats, Mann-Whitney test. (**B**) Heatmap of the immunoreactivity of anti-CV2 sera from male and female rats detected on a CRMP5 peptide array mapping the entire sequence of the protein in 15-mer peptide with 3 amino acid increments. (**C**) Bar graph with scatter plot showing the time to cross a sharp probe field in response to aversive (unsafe) bright light to reach the dark (safe) compartment. Rats immunized against CRMP5 and with validated mechanical hypersensitivity showed a longer time to cross the sharp probe field consistent with an aversive affective dimension of the probe stimulation. n=8 each group, *p<0.05, Kruskal-Wallis test. (**D**) Spinal cords from adult male and female rats injected with the CRMP5 coding plasmid compared to the control empty plasmid were used to assess potassium chloride (KCl, 90 mM)-induced CGRP release from nerve terminals. KCl triggered CGRP release which was significantly higher in spinal cords from rats with CRMP5 autoimmunity than in control rats (n=4 animals per group; *p<0.05, one-way ANOVA). Y-axis shows immunoreactive CGRP levels in the bath solution and normalized to the weight of each spinal cord.

**Supplementary Figure 7:**
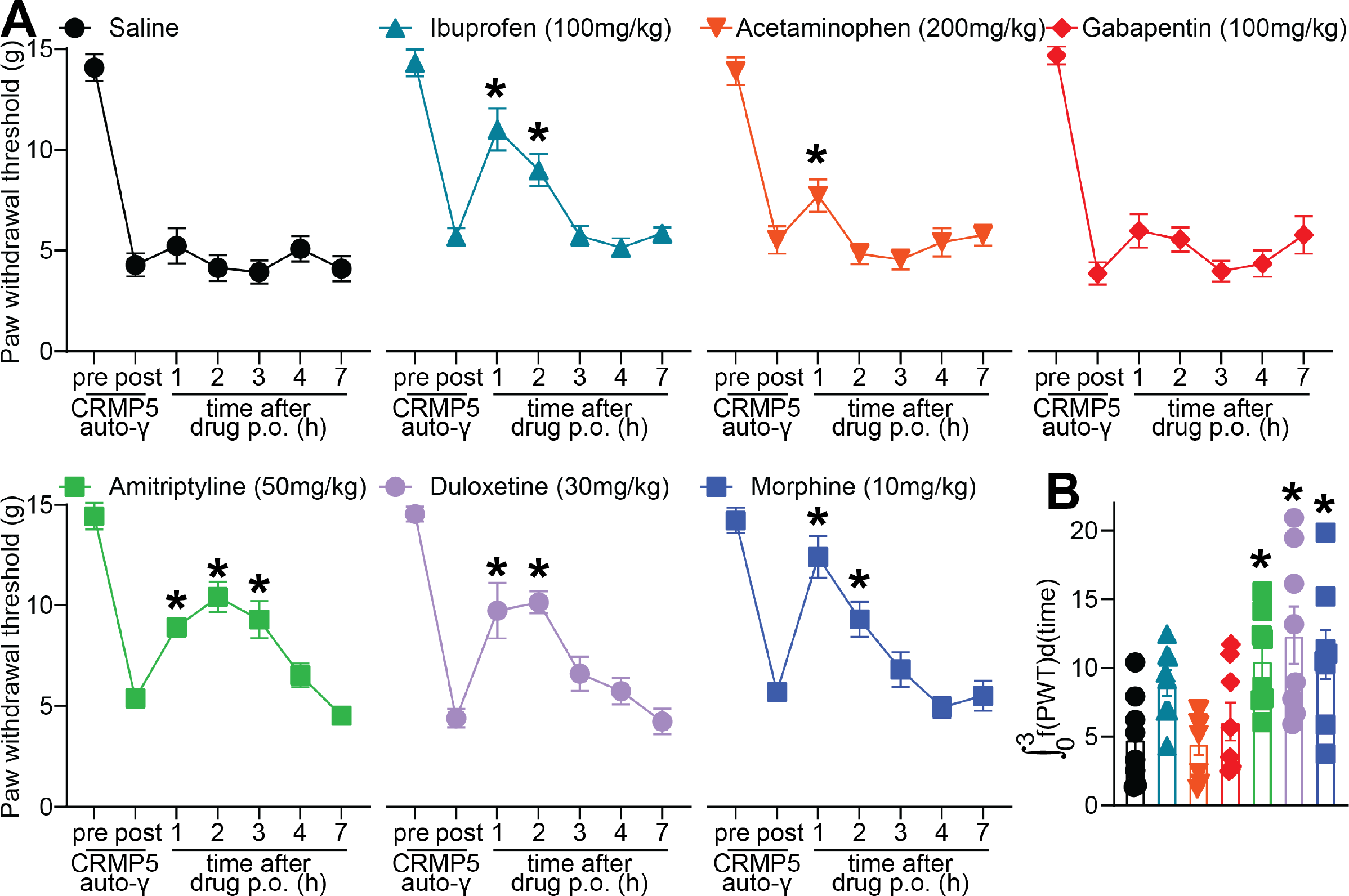
CRMP5 autoimmunity-induced mechanical hypersensitivity can be reversed by typical treatments for widespread pain. Rats with CRMP5 autoimmunity (auto-γ) and validated mechanical hypersensitivity received an oral administration of Ibuprofen (100mg/kg), acetaminophen (200mg/kg), gabapentin (100mg/kg), amitriptyline (50mg/kg), duloxetine (30mg/kg), or saline as a control. (**A**) Graphs showing the paw withdrawal threshold of rats treated as indicated. (**B**) Bar graph with scatter plot showing the curve integral (between 0 to 3 h) for each individual (n=8) from the indicated treatment groups. Lowered mechanical thresholds in rats with CRMP5 autoimmunity were reversed by amitriptyline, duloxetine and morphine while gabapentin had no effect. *p<0.05 compared to baseline. Data is shown as mean ± SEM. two-way ANOVA. Experimenters were blinded to groups and treatments.

**Supplementary Figure 8:**
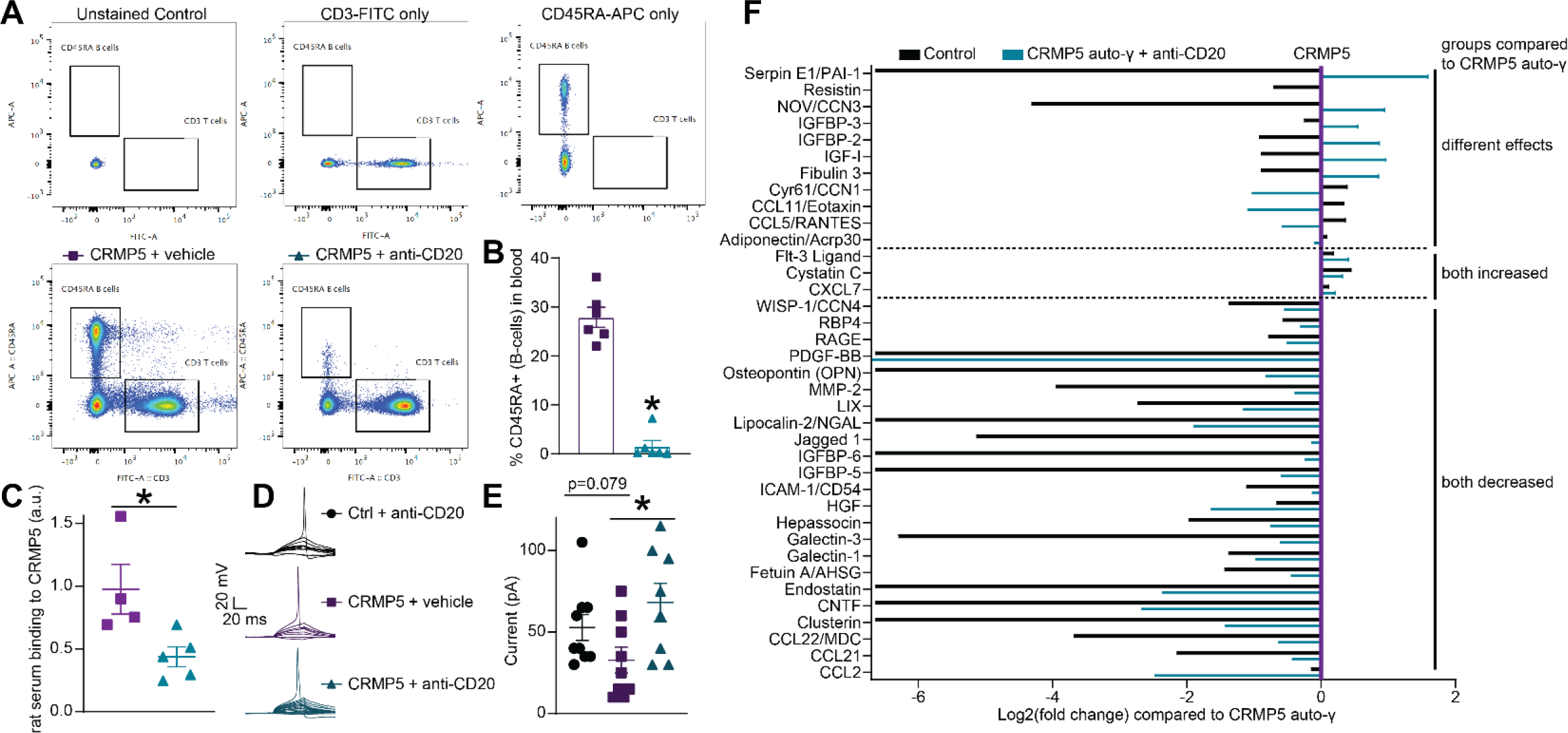
Anti-CD20 can deplete B-lymphocytes from rats with CRMP5 autoimmunity leading to increased rheobase in sensory neurons. (**A**) Gating method for detecting B- (CD45RA+) versus T- (CD3+) lymphocytes from rat blood, (**B**) Bar graph with scatter plot showing the percent of B-lymphocytes (CD45RA+) in the blood of rats with CRMP5 autoimmunity and injected with 1-mg of anti-CD20 as indicated. Anti-CD20 treatment efficiently depleted B-lymphocytes. (**C**) Scatter plot showing the level of rat serum from rats with CRMP5 autoimmunity and treated with anti-CD20 binding to CRMP5 in ELISA. Representative traces (**D**) and (**E**) bar graph with scatter plot showing the rheobase of sensory neurons cultured from rats with CRMP5 autoimmunity and treated with anti-CD20 as indicated. n=8-9 cells per condition; error bars indicate mean ± SEM, *p<0.05, Mann-Whitney test. (**F**) graph showing log2(fold change) of the indicated cytokines in the serum of control or treated with anti-CD20 rats normalized to serum from rats with CRMP5 autoimmunity (n=6 pooled).

**Table S1.** Details of statistical comparisons.

| Figure panel | Assay | Statistical Test; findings | Post-hoc analysis (adjusted p-values) | Number of subjects | Number of subjects excluded (ROUT test) |
| --- | --- | --- | --- | --- | --- |
| 1C | DRG neuron excitability, treated with #4 or #6 | Multiple Mann-Whitney tests | #4<br>current step (pA):<br>0 p= 0.214286<br>10 p= 0.191176<br>20 p= 0.027725<br>30 p= 0.025635<br>40 p= 0.031453<br>50 p= 0.002447<br>60 p= 0.004525<br>70 p= 0.008684<br>80 p= 0.004627<br>90 p= 0.004757<br>100 p= 0.006512<br>#6<br>current step (pA):<br>0 p= 0.47619<br>10 p= 0.148607<br>20 p= 0.058211<br>30 p= 0.007938<br>40 p= 0.00366<br>50 p= 0.001639<br>60 p= 0.001602<br>70 p= 0.004219<br>80 p= 0.004681<br>90 p= 0.007042<br>100 p= 0.005387 | PBS n= 11 cells<br>#4 n= 10 cells<br>#6 n= 11 cells | No exclusion |
| 1E | DRG neuron rheobase, treated with #4 or #6 | Kruskal-Wallis test | PBS vs #4<br>p= 0.0470<br>PBS vs #4<br>p= 0.0054 | PBS n= 14 cells<br>#4 n= 10 cells<br>#6 n= 12 cells | No exclusion |
| 1F | Anti-CV2 in rats i.th., paw withdrawal threshold | Kuskal-Wallis test<br>p<0.0001 | Dunn's multiple comparisons test, compared to baseline:<br>Serum with anti-CV2<br>1 h p= 0.0005<br>2 h p<0.0001<br>3 h p<0.0001<br>4 h p= 0.0001<br>5 h p= 0.0313 | Serum with anti-CV2: n= 3 rats per patients, 12 total | No exclusion |
| 1G | Anti-CV2 depleted serum in rats i.th., paw withdrawal threshold | Kuskal-Wallis test<br>P=0.0886 | Dunn's multiple comparisons test, compared to baseline:<br>CV2 depleted serum:<br>1 h p= 0.0515<br>2 h p= 0.0309<br>3 h p= 0.3017<br>4 h p= 0.1868<br>5 h p= 0.3659 | CV2 depleted serum: n= 3 rats per patients, 12 total | No exclusion |
| 1H | Anti-CV2 i.th. injection in rats, area under the curve | Mann-Whitney test | Anti-CV2 vs depleted serum<br>p<0.0001 | CV2 depleted serum: n= 3 rats per patients, 12 total | No exclusion |
| 2B | ELISA of epitope peptides blocking CV2/CRMP5 Abs binding to CRMP5 | One-way ANOVA | Patient #8 vs Patient #8 + peptides p=0.0004<br>Patient #8 vs Patient #8 + CRMP5 p=0.0004<br>Patient # 4 vs Patient #4 + peptides p<0.0001 | N= 3 replicates each group | No exclusion |
|  |  |  | Patient #4 vs Patient #4 + CRMP5 p=0.0002 |  |  |
| 2E | DRG neuron excitability, treated with patient #1 + blocking peptides | Two-way ANOVA, Current factor p<0.0001 Treatment factor p<0.0001 | Dunnett's multiple comparisons test<br>0 pA<br>0.03% DMSO vs. Patient #1 p= 0.9995<br>0.03% DMSO vs. Peptides p= 0.9995<br>0.03% DMSO vs. Patient #1 + Peptides p= 0.9995<br>10 pA<br>0.03% DMSO vs. Patient #1 p= 0.7479<br>0.03% DMSO vs. Peptides p= 0.9999<br>0.03% DMSO vs. Patient #1 + Peptides p=0.9864<br>20 pA<br>0.03% DMSO vs. Patient #1 p= 0.3546<br>0.03% DMSO vs. Peptides p>0.9999<br>0.03% DMSO vs. Patient #1 + Peptides p= 0.9959<br>30 pA<br>0.03% DMSO vs. Patient #1 p= 0.0459<br>0.03% DMSO vs. Peptides p= 0.9875<br>0.03% DMSO vs. Patient #1 + Peptides p= 0.9692<br>40 pA<br>0.03% DMSO vs. Patient #1 p= 0.0179<br>0.03% DMSO vs. Peptides p= 0.9796<br>0.03% DMSO vs. Patient #1 + Peptides p= 0.9959<br>50 pA<br>0.03% DMSO vs. Patient #1 p= 0.0082<br>0.03% DMSO vs. Peptides p= 0.9973<br>0.03% DMSO vs. Patient #1 + Peptides p>0.9999<br>60 pA<br>0.03% DMSO vs. Patient #1 p= 0.0038<br>0.03% DMSO vs. Peptides p= 0.9796<br>0.03% DMSO vs. Patient #1 + Peptides p= 0.9864<br>70 pA<br>0.03% DMSO vs. Patient #1 p= 0.0023<br>0.03% DMSO vs. Peptides p= 0.9005<br>0.03% DMSO vs. Patient #1 + Peptides p= 0.9429<br>80 pA | 0.03% DMSO n= 10 cells<br>Patient #1 n= 9 cells<br>0.03% DMSO + peptides n= 9 cells<br>Patient #1 + peptides | No exclusion |
|  |  |  | 0.03% DMSO vs. Patient #1 p= 0.0007<br>0.03% DMSO vs. Peptides p= 0.8665<br>0.03% DMSO vs. Patient #1 + Peptides p= 0.9429<br>90 pA<br>0.03% DMSO vs. Patient #1 p= 0.0006<br>0.03% DMSO vs. Peptides p= 0.9616<br>0.03% DMSO vs. Patient #1 + Peptides p= 0.9864<br>100 pA<br>0.03% DMSO vs. Patient #1 p= 0.0003<br>0.03% DMSO vs. Peptides p>0.9999<br>0.03% DMSO vs. Patient #1 + Peptides p= 0.9864 |  |  |
| 2G | DRG neuron Rheobase, treated with patient #1 + blocking peptides | Kuskal-Wallis test<br>P=0.4295 | Dunn's multiple comparisons test<br>Vehicle vs. 107-3 p>0.9999<br>Vehicle vs. Peptides p=0.9483<br>Vehicle vs. 107-3 + Peptides p>0.9999 | 0.03% DMSO n= 10 cells<br>Patient #1 n= 9 cells<br>0.03% DMSO + peptides n= 9 cells<br>Patient #1 + peptides | 1 cell excluded |
| 2I | Anti-CV2 in rats in paw, paw withdrawal threshold | Kuskal-Wallis test<br>p<0.0001 | Dunn's multiple comparisons test, compared to baseline:<br>Serum with anti-CV2<br>1 h p>0.9999<br>2 h p=0.8260<br>3 h p<0.0001<br>4 h p=0.0001<br>5 h p=0.0313 | Serum with anti-CV2 intraplantar: n= 3 rats per patients, 12 total | No exclusion |
| 2J | Anti-CV2 serum + peptides in rats in paw, paw withdrawal threshold | Kuskal-Wallis test<br>P=0.8368 | Dunn's multiple comparisons test, compared to baseline:<br>Anti-CV2 serum + peptides:<br>1 h p>0.9999<br>2 h p>0.9999<br>3 h p>0.9999<br>4 h p>0.9999<br>5 h p>0.9999 | Anti-CV2 serum + peptides: n= 3 rats per patients, 12 total | No exclusion |
| 2K | Anti-CV2 serum + peptides in paw, area under the curve | Mann-Whitney test | Anti-CV2 vs anti-CV2 + peptides<br>p<0.0001 | CV2 depleted serum: n= 3 rats per patients, 12 total | No exclusion |
| 3A | CRMP5 immunization, male rats, paw withdrawal threshold | Multiple Mann-Whitney tests | Control vs CRMP5 immunization at each time point (week):<br>0 p>0.999999<br>2 p>0.999999<br>4 p= 0.076923<br>6 p= 0.002642<br>8 p= 0.001399<br>10 p= 0.000155 | n= 8 rats each group | No exclusion |
| 3B | CRMP5 immunization, male rats, Area under the curve PWT | Mann-Whitney test | Control vs CRMP5 immunization:<br>p=0.0002 | n= 8 rats each group | No exclusion |
| 3C | CRMP5 immunization, male rats, paw withdrawal latency | Multiple Mann-Whitney tests | Control vs CRMP5 immunization at each time point (week):<br>0 p= 0.798446<br>2 p= 0.246465<br>4 p= 0.104895<br>6 p= 0.24359<br>8 p= 0.254079<br>10 p= 0.981818 | n= 8 rats each group | No exclusion |
| 3D | CRMP5 immunization, male rats, Area under the curve PWL | Mann-Whitney test | Control vs CRMP5 immunization:<br>p=0.9347 | n= 8 rats each group | No exclusion |
| 3E | CRMP5 immunization, female rats, paw withdrawal threshold | Multiple Mann-Whitney tests | Control vs CRMP5 immunization at each time point (week):<br>0 p>0.999999<br>2 p>0.999999<br>4 p= 0.006993<br>6 p= 0.000311<br>8 p= 0.001166<br>10 p= 0.000311 | n= 8 rats each group | No exclusion |
| 3F | CRMP5 immunization, female rats, Area under the curve PWT | Mann-Whitney test | Control vs CRMP5 immunization:<br>p=0.0002 | n= 8 rats each group | No exclusion |
| 3G | CRMP5 immunization, female rats, paw withdrawal latency | Multiple Mann-Whitney tests | Control vs CRMP5 immunization at each time point (week):<br>0 p= 0.382284<br>2 p= 0.289977<br>4 p= 0.858897<br>6 p= 0.625486<br>8 p= 0.093862<br>10 p= 0.441803 | n= 8 rats each group | No exclusion |
| 3H | CRMP5 immunization, female rats, Area under the curve PWL | Mann-Whitney test | Control vs CRMP5 immunization:<br>p=0.1304 | n= 8 rats each group | No exclusion |
| 3J | CRMP5 immunization, excitability | Multiple Mann-Whitney tests | Control vs CRMP5 immunization at each current step (pA):<br>Male<br>0 p>0.999999<br>10 p= 0.082836<br>20 p= 0.004268<br>30 p= 0.002417<br>40 p= 0.002674<br>50 p= 0.00617<br>60 p= 0.002005<br>70 p= 0.00036<br>80 p= 0.001337<br>90 p= 0.001028<br>Female<br>0 p>0.999999<br>10 p= 0.092066<br>20 p= 0.076036<br>30 p= 0.054517<br>40 p= 0.107143 | Male control n= 10<br>Male CRMP5 n=7<br>Female control n=11<br>Female control n=10 | No exclusion |
|  |  |  | 50 p= 0.046326<br>60 p= 0.013189<br>70 p= 0.009824<br>80 p= 0.01121<br>90 p= 0.009532 |  |  |
| 3L | CRMP5 immunization, rheobase | Mann-Whitney test | Control vs CRMP5 immunization:<br>Male p=0.0197<br>Female p=0.0521 | Male control n= 10<br>Male CRMP5 n=7<br>Female control n=11<br>Female control n=10 | No exclusion |
| 4A | CRMP5 autoimmunity treatment with Rituximab, withdrawal threshold | Two-way ANOVA<br>Time factor p<0.0001<br>Treatment factor p=0.0021 | CRMP5 autoimmunity treated with vehicle compared to pre-treatment baseline (D57):<br>day 57 vs day 60 p= 0.9997<br>day 57 vs day 64 p= 0.9748<br>day 57 vs day 67 p= 0.686<br>day 57 vs day 72 p= 0.7516<br>day 57 vs day 79 p= 0.998<br>day 57 vs day 82 p= 0.612<br>day 57 vs day 86 p>0.9999<br>day 57 vs day 60 p= 0.8809<br>CRMP5 autoimmunity treated with Rituximab compared to pre-treatment baseline (D57):<br>day 57 vs day 60 p= 0.9993<br>day 57 vs day 64 p= 0.7627<br>day 57 vs day 67 p= 0.9956<br>day 57 vs day 72 p= 0.2222<br>day 57 vs day 79 p= 0.016<br>day 57 vs day 82 p= 0.022<br>day 57 vs day 86 p= 0.0211<br>day 57 vs day 60 p= 0.0225 | Control + Anti-CD20 n=6<br>CRMP5 autoimmunity + Vehicle n=8<br>CRMP5 autoimmunity + anti-CD20 n=6 | No exclusion |
| 4B | CRMP5 autoimmunity treatment with Rituximab, area under the curve PWT | Mann-Whitney test | Vehicle vs anti-CD20 treatment: p=0.0007 | CRMP5 autoimmunity + Vehicle n=8<br>CRMP5 autoimmunity + anti-CD20 n=6 | No exclusion |
| 4D | CRMP5 autoimmunity treatment with Rituximab, excitability | Two-way ANOVA<br>Current factor p<0.0001<br>Treatment factor p<0.0001 | For each current step:<br>10 pA<br>Control + Anti-CD20 vs. CRMP5 autoimmunity + Vehicle p>0.9999<br>Control + Anti-CD20 vs. CRMP5 autoimmunity + Anti-CD20 p>0.9999<br>CRMP5 autoimmunity + Vehicle vs. CRMP5 autoimmunity + Anti-CD20 p>0.9999<br>20 pA<br>Control + Anti-CD20 vs. CRMP5 autoimmunity + Vehicle p= 0.7308<br>Control + Anti-CD20 vs. CRMP5 autoimmunity + Anti-CD20 p= 0.9855<br>CRMP5 autoimmunity + Vehicle vs. CRMP5 autoimmunity + Anti-CD20 p= 0.6437<br>30 pA | Control + Anti-CD20 n=9 cells<br>CRMP5 autoimmunity + Vehicle n=9 cells<br>CRMP5 autoimmunity + anti-CD20 n=8 cells | No exclusion |
|  |  |  | Control + Anti-CD20 vs. CRMP5 autoimmunity + Vehicle p= 0.373<br>Control + Anti-CD20 vs. CRMP5 autoimmunity + Anti-CD20 p= 0.9703<br>CRMP5 autoimmunity + Vehicle vs. CRMP5 autoimmunity + Anti-CD20 p= 0.2757<br>40 pA<br>Control + Anti-CD20 vs. CRMP5 autoimmunity + Vehicle p= 0.0923<br>Control + Anti-CD20 vs. CRMP5 autoimmunity + Anti-CD20 p= 0.9563<br>CRMP5 autoimmunity + Vehicle vs. CRMP5 autoimmunity + Anti-CD20 p= 0.0549<br>50 pA<br>Control + Anti-CD20 vs. CRMP5 autoimmunity + Vehicle p= 0.013<br>Control + Anti-CD20 vs. CRMP5 autoimmunity + Anti-CD20 p= 0.9931<br>CRMP5 autoimmunity + Vehicle vs. CRMP5 autoimmunity + Anti-CD20 p= 0.0228<br>60 pA<br>Control + Anti-CD20 vs. CRMP5 autoimmunity + Vehicle p= 0.0035<br>Control + Anti-CD20 vs. CRMP5 autoimmunity + Anti-CD20 p= 0.9931<br>CRMP5 autoimmunity + Vehicle vs. CRMP5 autoimmunity + Anti-CD20 p= 0.0033<br>70 pA<br>Control + Anti-CD20 vs. CRMP5 autoimmunity + Vehicle p= 0.0035<br>Control + Anti-CD20 vs. CRMP5 autoimmunity + Anti-CD20 p= 0.7861<br>CRMP5 autoimmunity + Vehicle vs. CRMP5 autoimmunity + Anti-CD20 p= 0.0005<br>80 pA<br>Control + Anti-CD20 vs. CRMP5 autoimmunity + Vehicle p= 0.0026<br>Control + Anti-CD20 vs. CRMP5 autoimmunity + Anti-CD20 p= 0.7565<br>CRMP5 autoimmunity + Vehicle vs. CRMP5 autoimmunity + Anti-CD20 p= 0.0003<br>90 pA |  |  |
| | | | Control + Anti-CD20 vs. CRMP5 autoimmunity + Vehicle $p=0.0008$<br>Control + Anti-CD20 vs. CRMP5 autoimmunity + Anti-CD20 $p=0.6693$<br>CRMP5 autoimmunity + Vehicle vs. CRMP5 autoimmunity + Anti-CD20 $p<0.0001$<br>100 pA<br>Control + Anti-CD20 vs. CRMP5 autoimmunity + Vehicle $p=0.0002$<br>Control + Anti-CD20 vs. CRMP5 autoimmunity + Anti-CD20 $p=0.831$<br>CRMP5 autoimmunity + Vehicle vs. CRMP5 autoimmunity + Anti-CD20 $p<0.0001$ | | |
| S6C | CRMP5 immunization, mechanical conflict avoidance | Multiple Mann-Whitney test | Male control vs Male CRMP5 $p=0.0499$<br>Female control vs Female CRMP5 $p=0.0002$ | Male control $n=8$<br>Male CRMP5 $n=8$<br>Female control $n=8$<br>Female CRMP5 $n=8$ | No exclusion |
| S6D | CRMP5 immunization, CGRP release | Two-way ANOVA<br>Fraction factor $p<0.0001$<br>Treatment factor $p=0.0051$ | Baseline 1<br>Male control vs. Male CRMP5 $p=0.8599$<br>Male control vs. Female control $p=0.3253$<br>Male control vs. Female CRMP5 $p=0.896$<br>Male CRMP5 vs. Female control $p=0.0789$<br>Male CRMP5 vs. Female CRMP5 $p=0.9961$<br>Female control vs. Female CRMP5 $p=0.3356$<br>Baseline 2<br>Male control vs. Male CRMP5 $p=0.414$<br>Male control vs. Female control $p=0.3049$<br>Male control vs. Female CRMP5 $p=0.7755$<br>Male CRMP5 vs. Female control $p=0.1042$<br>Male CRMP5 vs. Female CRMP5 $p=0.9992$<br>Female control vs. Female CRMP5 $p=0.3824$<br>90 mM KCl<br>Male control vs. Male CRMP5 $p=0.0486$<br>Male control vs. Female control $p=0.9997$<br>Male control vs. Female CRMP5 $p=0.0586$<br>Male CRMP5 vs. Female control $p=0.0118$ | Male control $n=4$<br>Male CRMP5 $n=4$<br>Female control $n=4$<br>Female CRMP5 $n=4$ | No exclusion |
|  |  |  | Male CRMP5 vs. Female CRMP5 p= 0.9958<br>Female control vs. Female CRMP5 p= 0.0401<br>Wash 1<br>Male control vs. Male CRMP5 p= 0.5929<br>Male control vs. Female control p= 0.4595<br>Male control vs. Female CRMP5 p= 0.9951<br>Male CRMP5 vs. Female control p= 0.1902<br>Male CRMP5 vs. Female CRMP5 p= 0.5466<br>Female control vs. Female CRMP5 p= 0.7099<br>Wash 2<br>Male control vs. Male CRMP5 p= 0.962<br>Male control vs. Female control p= 0.3584<br>Male control vs. Female CRMP5 p= 0.6683<br>Male CRMP5 vs. Female control p= 0.2047<br>Male CRMP5 vs. Female CRMP5 p= 0.3961<br>Female control vs. Female CRMP5 p= 0.1528 |  |  |
| S7A | CRMP5 autoimmunity treatment with painkillers, withdrawal threshold | Two-way ANOVA<br>Time factor p<0.0001<br>Treatment factor p=0.0051 | Saline<br>Post vs. Pre p= 0.0002<br>Post vs. 1h p= 0.863<br>Post vs. 2h p>0.9999<br>Post vs. 4h p= 0.994<br>Post vs. 5h p= 0.929<br>Post vs. 7h p= 0.9999<br>Morphine (10mg/kg)<br>Post vs. Pre p<0.0001<br>Post vs. 1h p= 0.0018<br>Post vs. 2h p= 0.0186<br>Post vs. 4h p= 0.5014<br>Post vs. 5h p= 0.6986<br>Post vs. 7h p= 0.9996<br>Ibuprofen (100mg/kg)<br>Post vs. Pre p= 0.0001<br>Post vs. 1h p= 0.0021<br>Post vs. 2h p= 0.0124<br>Post vs. 4h p>0.9999<br>Post vs. 5h p= 0.8186<br>Post vs. 7h p= 0.991<br>Acetaminophen (200mg/kg)<br>Post vs. Pre p<0.0001<br>Post vs. 1h p= 0.0199<br>Post vs. 2h p= 0.939<br>Post vs. 4h p= 0.6289<br>Post vs. 5h p>0.9999<br>Post vs. 7h p= 0.9963<br>Gabapentin (100mg/kg)<br>Post vs. Pre p<0.0001 | Saline n=8<br>Morphine (10mg/kg) n=8<br>Ibuprofen (100mg/kg) n=8<br>Acetaminophen (200mg/kg) n=8<br>Gabapentin (100mg/kg) n=8<br>Duloxetine (30mg/kg) n=8<br>Amitriptyline (50mg/kg) n=8 | No exclusion |
|  |  |  | Post vs. 1h p= 0.3542<br>Post vs. 2h p= 0.3569<br>Post vs. 4h p>0.9999<br>Post vs. 5h p= 0.9884<br>Post vs. 7h p= 0.1402<br>Duloxetine (30mg/kg)<br>Post vs. Pre p<0.0001<br>Post vs. 1h p= 0.0468<br>Post vs. 2h p= 0.0005<br>Post vs. 4h p= 0.1432<br>Post vs. 5h p= 0.1287<br>Post vs. 7h p>0.9999<br>Amitriptyline (50mg/kg)<br>Post vs. Pre p<0.0001<br>Post vs. 1h p= 0.0017<br>Post vs. 2h p= 0.0011<br>Post vs. 4h p= 0.0384<br>Post vs. 5h p= 0.65<br>Post vs. 7h p= 0.4698 |  |  |
| S7B | CRMP5 autoimmunity treatment with painkillers, area under the curve | One way ANOVA with Holm-Šídák's multiple comparisons test | Saline vs. Morphine (10mg/kg) p= 0.0161<br>Saline vs. Ibuprofen (100mg/kg) p= 0.1309<br>Saline vs. Acetaminophen (200mg/kg) p= 0.878<br>Saline vs. Gabapentin (100mg/kg) p= 0.7724<br>Saline vs. Duloxetine (30mg/kg) p= 0.0024<br>Saline vs. Amitriptyline (50mg/kg) p= 0.0237 | Saline n=8<br>Morphine (10mg/kg) n=8<br>Ibuprofen (100mg/kg) n=8<br>Acetaminophen (200mg/kg) n=8<br>Gabapentin (100mg/kg) n=8<br>Duloxetine (30mg/kg) n=8<br>Amitriptyline (50mg/kg) n=8 | No exclusion |
| S8B | CRMP5 autoimmunity treatment with Rituximab, %CD45RA+ cells | Mann-Whitney test | CRMP5 autoimmunity + Vehicle vs. CRMP5 autoimmunity + Anti-CD20 p=0.0022 | CRMP5 autoimmunity + Vehicle n=6<br>CRMP5 autoimmunity + anti-CD20 n=6 | No exclusion |
| S8B | CRMP5 autoimmunity treatment with Rituximab, binding to CRMP5 | Mann-Whitney test | CRMP5 autoimmunity + Vehicle vs. CRMP5 autoimmunity + Anti-CD20 p=0.0238 | CRMP5 autoimmunity + Vehicle n=4<br>CRMP5 autoimmunity + anti-CD20 n=6 | No exclusion |
| S8E | CRMP5 autoimmunity treatment with Rituximab, Rheobase | Multiple Mann-Whitney tests | Control + Anti-CD20 vs. CRMP5 autoimmunity + Vehicle p=0.0799<br>CRMP5 autoimmunity + Vehicle vs. CRMP5 autoimmunity + Anti-CD20 p=0.0248 | Control + Anti-CD20 n=9 cells<br>CRMP5 autoimmunity + Vehicle n=9 cells<br>CRMP5 autoimmunity + anti-CD20 n=8 cells | No exclusion |

